# Broken, silent, and in hiding: Tamed endogenous pararetroviruses escape elimination from the genome of sugar beet (*Beta vulgaris*)

**DOI:** 10.1101/2020.12.02.407783

**Authors:** Nicola Schmidt, Kathrin M. Seibt, Beatrice Weber, Trude Schwarzacher, Thomas Schmidt, Tony Heitkam

**Affiliations:** Institute of Botany, Technische Universität Dresden, 01069 Dresden, Germany; Department of Genetics and Genome Biology, University of Leicester, LE1 7RH Leicester, United Kingdom; Key Laboratory of Plant Resources Conservation and Sustainable Utilization / Guangdong Provincial Key Laboratory of Applied Botany, South China Botanical Garden, Chinese Academy of Sciences, Xingke Road 723, Tianhe District, Guangzhou, 510650, PR China

**Keywords:** *Beta vulgaris*, sugar beet, pararetrovirus, *Caulimoviridae*, *Florendovirus*, retrotransposon, fluorescent *in situ* hybridization

## Abstract

**Background and Aims:** Endogenous pararetroviruses (EPRVs) are widespread components of plant genomes that originated from episomal DNA viruses of the *Caulimoviridae* family. Due to fragmentation and rearrangements, most EPRVs have lost their ability to replicate through reverse transcription and to initiate viral infection. Similar to the closely related retrotransposons, extant EPRVs were retained and often amplified in plant genomes for several million years. Here, we characterize the complete genomic EPRV fraction of the crop sugar beet (*Beta vulgaris*, Amaranthaceae) to understand how they shaped the beet genome and to suggest explanations for their absent virulence.

**Methods:** Using next- and third-generation sequencing data and the genome assembly, we reconstructed full-length *in silico* representatives for the three host-specific EPRV families (beetEPRVs) in the *B. vulgaris* genome. Focusing on the canonical family beetEPRV3, we investigated its chromosomal localization, abundance, and distribution by fluorescent *in situ* and Southern hybridization.

**Key Results:** BeetEPRVs range between 7.5 and 10.7 kb (0.3 % of the *B. vulgaris* genome) and are heterogeneous in structure and sequence. Although all three beetEPRV families were assigned to the florendoviruses, they showed variably arranged protein-coding domains, different degrees of fragmentation, and preferences for diverse sequence contexts. We observed small RNAs that target beetEPRVs in a family-specific manner, indicating stringent epigenetic suppression. We localized beetEPRV3 on all 18 sugar beet chromosomes, occurring preferentially in clusters and associated with heterochromatic, centromeric and intercalary satellite DNAs. BeetEPRV3 variants also exist in the genomes of related wild species, indicating an initial beetEPRV3 integration 13.4 to 7.2 million years ago.

**Conclusions:** Our study in beet illustrates the variability of EPRV structure and sequence in a single host genome. Evidence of sequence fragmentation and epigenetic silencing imply possible plant strategies to cope with long-term persistence of EPRVs, including amplification, fixation in the heterochromatin, and containment of EPRV virulence.

## INTRODUCTION

Endogenous pararetroviruses (EPRVs) are viral double-stranded nucleic acids that permanently reside in the genome of their host. In plants, the ancestral EPRV progenitors are exogenous viruses of the *Caulimoviridae* family that integrated into the host nuclear genome through illegitimate recombination several million years ago (Jakowitsch *et al.*, 1999; Diop *et al.*, 2018). Caulimoviruses are reverse-transcribing viruses (*Ortervirales*; reviewed by Krupovic *et al.*, 2018; Teycheney *et al.*, 2020); though, in contrast to the closely related retroviruses (e.g. HIV; International Committee on Taxonomy of Viruses, 2019), integration into the host genome is not obligatory for their replication. Over time, nearly all EPRVs underwent fragmentation and rearrangements within the host genome, thus losing their activity. However, in some cases, these ancient integrated viral sequences can be activated through reverse transcription and recombination, thereby forming virulent episomes often associated with devastating diseases (reviewed by Staginnus and Richert-Pöggeler, 2006; Chabannes and Iskra-Caruana, 2013; Kuriyama *et al.*, 2020).

As EPRVs exist in a broad range of vascular plants (Diop *et al.*, 2018; Gong and Han, 2018) and cover a wider spectrum of genera than the exogenous *Caulimoviridae*, an extinction of several homologous episomal counterparts is repeatedly assumed (reviewed by Chen and Kishima, 2016). In general, the caulimoviral genera are characterized by differences in their nucleotide sequence and the organization of the viral genome, e.g. the number of open reading frames (ORFs) and the arrangement of essential protein domains within them. For instance, petuviruses typically have one single ORF (Richert-Pöggeler *et al.*, 2003), whereas caulimoviruses and solendoviruses differ in the allocation of the protein domains in their four ORFs (Geering *et al.*, 2010). Using gene order and ORF arrangements, further genera were defined: Among those, the florendoviruses (FEVs) encode the characteristic movement protein (MP), the coat protein with a zinc finger motif (ZF), the aspartic protease (AP), the reverse trancriptase (RT), and the ribonuclease H1 (RH) on the first of two overlapping ORFs (Geering *et al.*, 2014). The florendoviruses are among the most abundant EPRVs in the plant kingdom, occurring in economically important plants such as *Elaeis guineensis* (oil palm), *Gossypium raimondii* (cotton), *Citrus* × *sinensis* (orange), *Glycine max* (soybean), *Petunia* sp. and *Beta vulgaris* (sugar beet; Bombarely *et al.*, 2016; Diop *et al.*, 2018). Although pararetroviruses have been detected in many plant genomes, it has not yet been possible to resolve their complex organization in the deep AT-rich heterochromatin.

In sugar beet, pararetroviruses contribute about 0.4-0.5 % to the genome (Dohm *et al.*, 2014; Diop *et al.*, 2018). As we do not know of any outbreaks of associated diseases, beet’s endogenous viral sequences have likely been assimilated by the host. Therefore, beet may represent a suitable organism to study how the host genome buries, disassembles and inactivates potentially destructive sequences.

Sugar beet is one of the most important crops of the moderate climate zones, contributing to approximately 14 % of the world’s sugar production (FAOSTAT, 2017). Cultivated beet species and related wild beets belong to the sister genera *Beta* and *Patellifolia* within the Amaranthaceae. According to Ulbrich (1934) and Frese *et al.* (2000), the genus *Beta* can be further subdivided into the three sections *Beta*, *Corollinae*, and *Nanae*. A comparison to wild beet genomes may offer an insight into the acquisition of EPRVs in the *Beta* genus.

Reference genome sequences and long read information are already available for two *. vulgaris* genotypes (Dohm *et al.*, 2014; Funk *et al.*, 2018; McGrath *et al.*, 2020). Similar to the euchromatic genic regions of beet, its heterochromatin is well-studied (Schmidt and Heslop-Harrison, 1998) and consists to a large part of satellite DNA (satDNA), such as the centromeric satDNA family pBV (Schmidt and Metzlaff, 1991; Zakrzewski *et al.*, 2013) and the intercalary satDNA family pEV (Schmidt *et al.*, 1991). Here, a combination of bioinformatics, advanced genomics, and molecular cytogenetics is used to investigate how the genome of beet may repress and disassemble EPRVs. For this, we characterize the EPRV landscape in the sugar beet genome and resolve the highly repetitive environment. Finally, we test if beetEPRVs are target for silencing by small RNAs, presumably involved in protecting the genome from subsequent beetEPRV infection. Thus, we aim to illustrate the variety EPRVs can attain in a single host and within the same genus, and we provide a possible explanation for the ability of EPRVs to escape elimination after ancient infection events – by integration into preserving genomic environments and by contribution to the host’s defense against EPRV-derived pathogens.

## MATERIALS AND METHODS

### Bioinformatic identification of B. vulgaris-specific EPRVs

To enable a targeted EPRV detection, we collected thirteen publicly available EPRVs, including the nine sequences from *gydb.org* (Llorens *et al.*, 2009; Llorens *et al.*, 2011), FriEPRV (Becher *et al.*, 2014), and the florendoviruses *Atrich*BV, *Gmax*V, and *Ljap*AV (Geering *et al.*, 2014). Subsequently, representative sequences of beetEPRV1, beetEPRV2, beetEPRV3 were added (Supplementary data S1-S5). The EPRV reference set therefore contained 16 sequences in total, representing the EPRV genera *Petu-*, *Badna-*, *Caulimo-*, *Cavemo-*, *Solendo*-, *Soymo-, Tungrovirus*, and *Florendovirus* (Table S1). After identification of their MP and RT domains, we aligned the respective nucleic acid sequences using *MAFFT* (Katoh and Standley, 2013) followed by manual refinement to build nucleotide Hidden Markov Models (nHMMs).

Using these nHMMs with *nhmmer* (Wheeler and Eddy, 2013), we identified the EPRV RT and MP sequences from the sugar beet EL10.1 assembly (Funk *et al.*, 2018; McGrath *et al.*, 2020) as well as the corresponding single molecule real-time (SMRT) reads (genebank accession SRX3402137). The results were parsed to analyze the hits, choose cut-off parameters and extract the corresponding sequences. After parameter analysis (Supplementary data Fig. S1), we selected all detected 262 (assembly) and 350 (SMRT reads) MP hits for further analysis. In contrast, to avoid cross-detection of similar Ty3-*gypsy* RTs, 125 assembly-derived and 320 SMRT read-derived RT sequences with an nHMM coverage of at least 200 bp and an nHMM coverage/bitscore quotient between 1.5 and 2.5 have been considered. From the assembly RT hits, another six candidates were excluded: five showed a high degree of fragmentation, and one represented a Ty3-*gypsy* sequence. Therefore, 119 assembly RT hits remained for our analysis. Visualizations of the *nhmmer* search results were created using Python v. 2.7 with the seaborn package (Waskom *et al.*, 2018).

To identify potentially intact members of the beetEPRV families, we screened the flanking region (± 8 kb) of the beetEPRV RTs for adjacent MP domains, subsequently excluding RT fragments.

### Sequence analyses and comparisons

Multiple sequence alignments have been calculated with *MUSCLE* and *MAFFT* (Edgar, 2004; Katoh and Standley, 2013), followed by manual refinement. To generate representative reference sequences for each beetEPRV family (Supplementary data S1), we have built consensuses from alignments of 11 to 42 sequences, respectively (Supplementary data S2-5; Table 1). Secondary structures for the beetEPRV consensus elements were predicted with JPred 4.0 (Drozdetskiy *et al.*, 2015). The presumed weights of the encoded proteins were determined by the Protein Molecular Weight Calculator (sciencegateway.org/tools/proteinmw.htm).

**Table 1:**
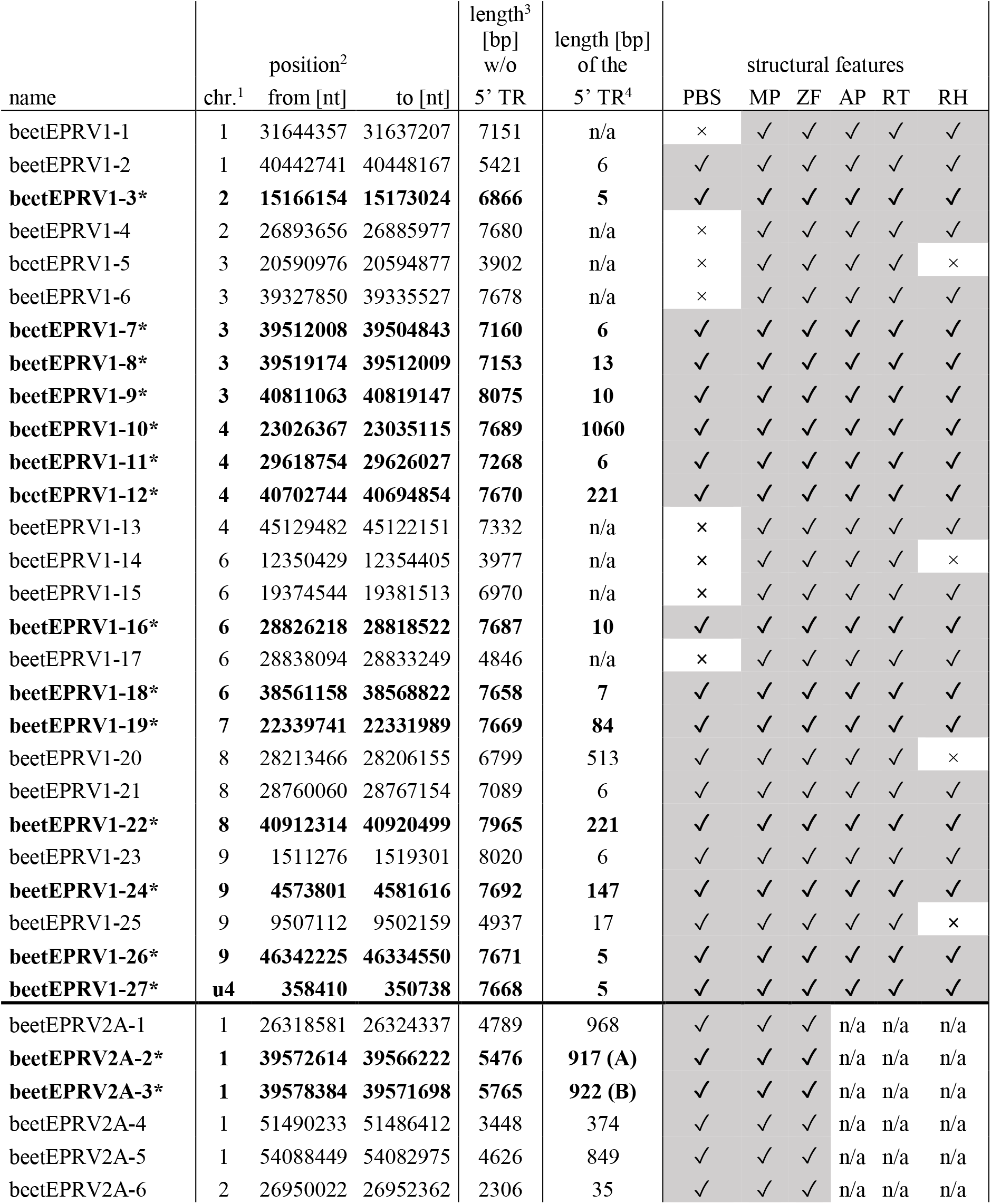
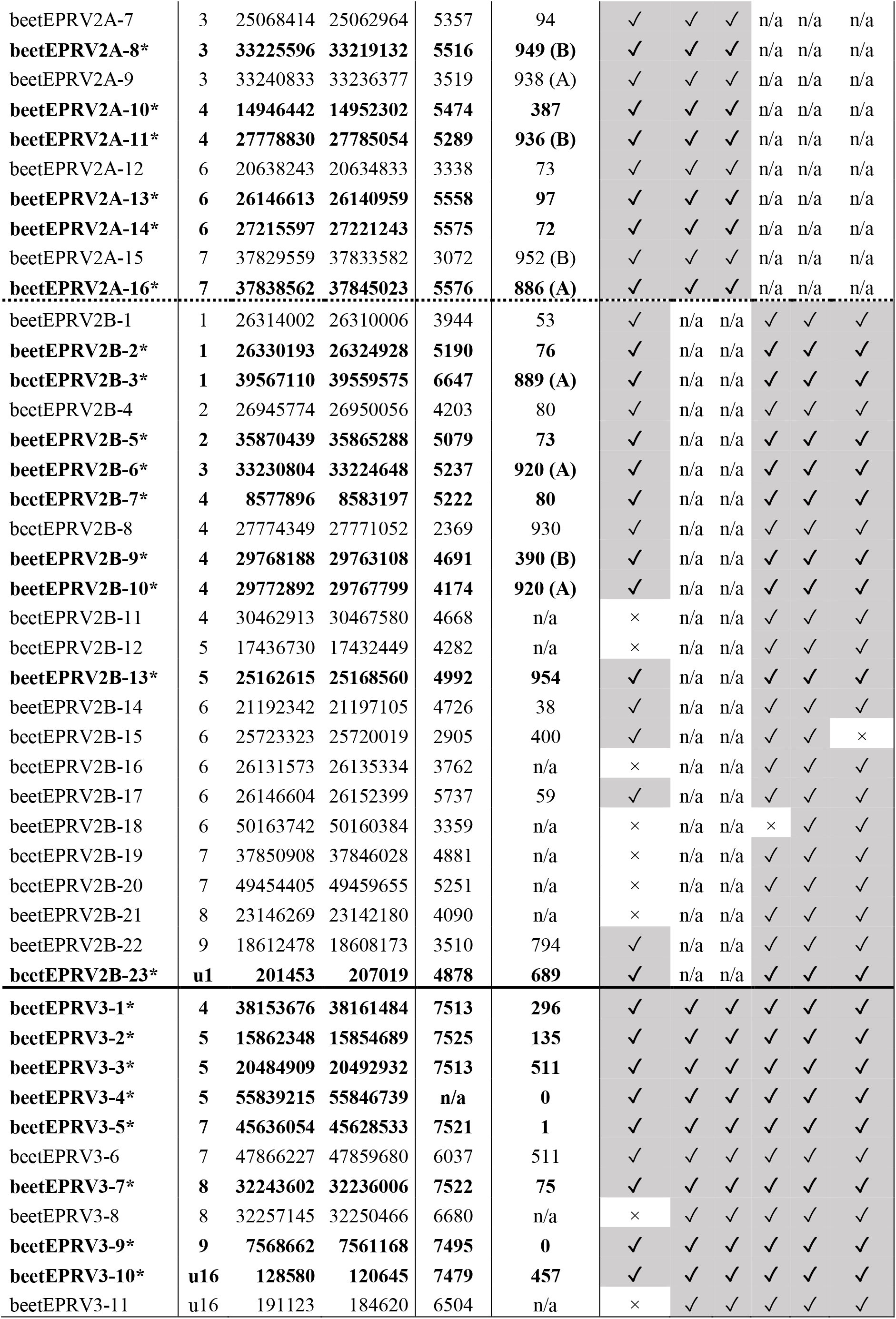
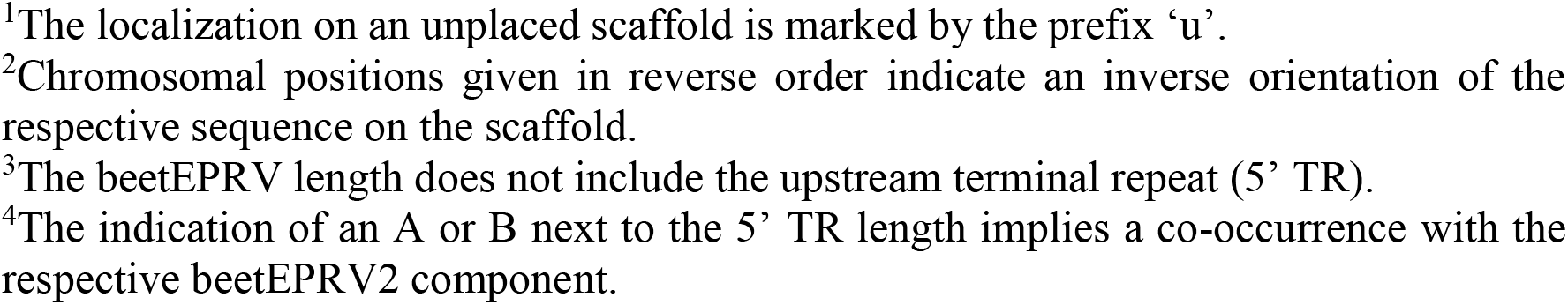
Chromosomal position of beetEPRV sequences (found via *nhmmer* in the EL10 assembly; Funk *et al.*, 2018) sorted by family. Asterisks behind the sequence name (bold entries) indicate canonical full-length beetEPRV sequences according to the presence of all protein domains (check mark in every “structural features” column) and the completeness of the ORF(s) (bold check marks).

| name | position<sup>2</sup> |  |  | length<sup>3</sup><br>[bp]<br>w/o | length [bp]<br>of the | structural features |  |  |  |  |  |
| --- | --- | --- | --- | --- | --- | --- | --- | --- | --- | --- | --- |
|  | chr.<sup>1</sup> | from [nt] | to [nt] | 5' TR | 5' TR<sup>4</sup> | PBS | MP | ZF | AP | RT | RH |
| beetEPRV1-1 | 1 | 31644357 | 31637207 | 7151 | n/a | × | ✓ | ✓ | ✓ | ✓ | ✓ |
| beetEPRV1-2 | 1 | 40442741 | 40448167 | 5421 | 6 | ✓ | ✓ | ✓ | ✓ | ✓ | ✓ |
| **beetEPRV1-3\*** | **2** | **15166154** | **15173024** | **6866** | **5** | ✓ | ✓ | ✓ | ✓ | ✓ | ✓ |
| beetEPRV1-4 | 2 | 26893656 | 26885977 | 7680 | n/a | × | ✓ | ✓ | ✓ | ✓ | ✓ |
| beetEPRV1-5 | 3 | 20590976 | 20594877 | 3902 | n/a | × | ✓ | ✓ | ✓ | ✓ | × |
| beetEPRV1-6 | 3 | 39327850 | 39335527 | 7678 | n/a | × | ✓ | ✓ | ✓ | ✓ | ✓ |
| **beetEPRV1-7\*** | **3** | **39512008** | **39504843** | **7160** | **6** | ✓ | ✓ | ✓ | ✓ | ✓ | ✓ |
| **beetEPRV1-8\*** | **3** | **39519174** | **39512009** | **7153** | **13** | ✓ | ✓ | ✓ | ✓ | ✓ | ✓ |
| **beetEPRV1-9\*** | **3** | **40811063** | **40819147** | **8075** | **10** | ✓ | ✓ | ✓ | ✓ | ✓ | ✓ |
| **beetEPRV1-10\*** | **4** | **23026367** | **23035115** | **7689** | **1060** | ✓ | ✓ | ✓ | ✓ | ✓ | ✓ |
| **beetEPRV1-11\*** | **4** | **29618754** | **29626027** | **7268** | **6** | ✓ | ✓ | ✓ | ✓ | ✓ | ✓ |
| **beetEPRV1-12\*** | **4** | **40702744** | **40694854** | **7670** | **221** | ✓ | ✓ | ✓ | ✓ | ✓ | ✓ |
| beetEPRV1-13 | 4 | 45129482 | 45122151 | 7332 | n/a | × | ✓ | ✓ | ✓ | ✓ | ✓ |
| beetEPRV1-14 | 6 | 12350429 | 12354405 | 3977 | n/a | × | ✓ | ✓ | ✓ | ✓ | × |
| beetEPRV1-15 | 6 | 19374544 | 19381513 | 6970 | n/a | × | ✓ | ✓ | ✓ | ✓ | ✓ |
| **beetEPRV1-16\*** | **6** | **28826218** | **28818522** | **7687** | **10** | ✓ | ✓ | ✓ | ✓ | ✓ | ✓ |
| beetEPRV1-17 | 6 | 28838094 | 28833249 | 4846 | n/a | × | ✓ | ✓ | ✓ | ✓ | ✓ |
| **beetEPRV1-18\*** | **6** | **38561158** | **38568822** | **7658** | **7** | ✓ | ✓ | ✓ | ✓ | ✓ | ✓ |
| **beetEPRV1-19\*** | **7** | **22339741** | **22331989** | **7669** | **84** | ✓ | ✓ | ✓ | ✓ | ✓ | ✓ |
| beetEPRV1-20 | 8 | 28213466 | 28206155 | 6799 | 513 | ✓ | ✓ | ✓ | ✓ | ✓ | × |
| beetEPRV1-21 | 8 | 28760060 | 28767154 | 7089 | 6 | ✓ | ✓ | ✓ | ✓ | ✓ | ✓ |
| **beetEPRV1-22\*** | **8** | **40912314** | **40920499** | **7965** | **221** | ✓ | ✓ | ✓ | ✓ | ✓ | ✓ |
| beetEPRV1-23 | 9 | 1511276 | 1519301 | 8020 | 6 | ✓ | ✓ | ✓ | ✓ | ✓ | ✓ |
| **beetEPRV1-24\*** | **9** | **4573801** | **4581616** | **7692** | **147** | ✓ | ✓ | ✓ | ✓ | ✓ | ✓ |
| beetEPRV1-25 | 9 | 9507112 | 9502159 | 4937 | 17 | ✓ | ✓ | ✓ | ✓ | ✓ | × |
| **beetEPRV1-26\*** | **9** | **46342225** | **46334550** | **7671** | **5** | ✓ | ✓ | ✓ | ✓ | ✓ | ✓ |
| **beetEPRV1-27\*** | **u4** | **358410** | **350738** | **7668** | **5** | ✓ | ✓ | ✓ | ✓ | ✓ | ✓ |
| beetEPRV2A-1 | 1 | 26318581 | 26324337 | 4789 | 968 | ✓ | ✓ | ✓ | n/a | n/a | n/a |
| **beetEPRV2A-2\*** | **1** | **39572614** | **39566222** | **5476** | **917 (A)** | ✓ | ✓ | ✓ | n/a | n/a | n/a |
| **beetEPRV2A-3\*** | **1** | **39578384** | **39571698** | **5765** | **922 (B)** | ✓ | ✓ | ✓ | n/a | n/a | n/a |
| beetEPRV2A-4 | 1 | 51490233 | 51486412 | 3448 | 374 | ✓ | ✓ | ✓ | n/a | n/a | n/a |
| beetEPRV2A-5 | 1 | 54088449 | 54082975 | 4626 | 849 | ✓ | ✓ | ✓ | n/a | n/a | n/a |
| beetEPRV2A-6 | 2 | 26950022 | 26952362 | 2306 | 35 | ✓ | ✓ | ✓ | n/a | n/a | n/a |
| beetEPRV2A-7 | 3 | 25068414 | 25062964 | 5357 | 94 | ✓ | ✓ | ✓ | n/a | n/a | n/a |
| **beetEPRV2A-8\*** | **3** | **33225596** | **33219132** | **5516** | **949 (B)** | ✓ | ✓ | ✓ | n/a | n/a | n/a |
| beetEPRV2A-9 | 3 | 33240833 | 33236377 | 3519 | 938 (A) | ✓ | ✓ | ✓ | n/a | n/a | n/a |
| **beetEPRV2A-10\*** | **4** | **14946442** | **14952302** | **5474** | **387** | ✓ | ✓ | ✓ | n/a | n/a | n/a |
| **beetEPRV2A-11\*** | **4** | **27778830** | **27785054** | **5289** | **936 (B)** | ✓ | ✓ | ✓ | n/a | n/a | n/a |
| beetEPRV2A-12 | 6 | 20638243 | 20634833 | 3338 | 73 | ✓ | ✓ | ✓ | n/a | n/a | n/a |
| **beetEPRV2A-13\*** | **6** | **26146613** | **26140959** | **5558** | **97** | ✓ | ✓ | ✓ | n/a | n/a | n/a |
| **beetEPRV2A-14\*** | **6** | **27215597** | **27221243** | **5575** | **72** | ✓ | ✓ | ✓ | n/a | n/a | n/a |
| beetEPRV2A-15 | 7 | 37829559 | 37833582 | 3072 | 952 (B) | ✓ | ✓ | ✓ | n/a | n/a | n/a |
| **beetEPRV2A-16\*** | **7** | **37838562** | **37845023** | **5576** | **886 (A)** | ✓ | ✓ | ✓ | n/a | n/a | n/a |
| beetEPRV2B-1 | 1 | 26314002 | 26310006 | 3944 | 53 | ✓ | n/a | n/a | ✓ | ✓ | ✓ |
| **beetEPRV2B-2\*** | **1** | **26330193** | **26324928** | **5190** | **76** | ✓ | n/a | n/a | ✓ | ✓ | ✓ |
| **beetEPRV2B-3\*** | **1** | **39567110** | **39559575** | **6647** | **889 (A)** | ✓ | n/a | n/a | ✓ | ✓ | ✓ |
| beetEPRV2B-4 | 2 | 26945774 | 26950056 | 4203 | 80 | ✓ | n/a | n/a | ✓ | ✓ | ✓ |
| **beetEPRV2B-5\*** | **2** | **35870439** | **35865288** | **5079** | **73** | ✓ | n/a | n/a | ✓ | ✓ | ✓ |
| **beetEPRV2B-6\*** | **3** | **33230804** | **33224648** | **5237** | **920 (A)** | ✓ | n/a | n/a | ✓ | ✓ | ✓ |
| **beetEPRV2B-7\*** | **4** | **8577896** | **8583197** | **5222** | **80** | ✓ | n/a | n/a | ✓ | ✓ | ✓ |
| beetEPRV2B-8 | 4 | 27774349 | 27771052 | 2369 | 930 | ✓ | n/a | n/a | ✓ | ✓ | ✓ |
| **beetEPRV2B-9\*** | **4** | **29768188** | **29763108** | **4691** | **390 (B)** | ✓ | n/a | n/a | ✓ | ✓ | ✓ |
| **beetEPRV2B-10\*** | **4** | **29772892** | **29767799** | **4174** | **920 (A)** | ✓ | n/a | n/a | ✓ | ✓ | ✓ |
| beetEPRV2B-11 | 4 | 30462913 | 30467580 | 4668 | n/a | × | n/a | n/a | ✓ | ✓ | ✓ |
| beetEPRV2B-12 | 5 | 17436730 | 17432449 | 4282 | n/a | × | n/a | n/a | ✓ | ✓ | ✓ |
| **beetEPRV2B-13\*** | **5** | **25162615** | **25168560** | **4992** | **954** | ✓ | n/a | n/a | ✓ | ✓ | ✓ |
| beetEPRV2B-14 | 6 | 21192342 | 21197105 | 4726 | 38 | ✓ | n/a | n/a | ✓ | ✓ | ✓ |
| beetEPRV2B-15 | 6 | 25723323 | 25720019 | 2905 | 400 | ✓ | n/a | n/a | ✓ | ✓ | × |
| beetEPRV2B-16 | 6 | 26131573 | 26135334 | 3762 | n/a | × | n/a | n/a | ✓ | ✓ | ✓ |
| beetEPRV2B-17 | 6 | 26146604 | 26152399 | 5737 | 59 | ✓ | n/a | n/a | ✓ | ✓ | ✓ |
| beetEPRV2B-18 | 6 | 50163742 | 50160384 | 3359 | n/a | × | n/a | n/a | × | ✓ | ✓ |
| beetEPRV2B-19 | 7 | 37850908 | 37846028 | 4881 | n/a | × | n/a | n/a | ✓ | ✓ | ✓ |
| beetEPRV2B-20 | 7 | 49454405 | 49459655 | 5251 | n/a | × | n/a | n/a | ✓ | ✓ | ✓ |
| beetEPRV2B-21 | 8 | 23146269 | 23142180 | 4090 | n/a | × | n/a | n/a | ✓ | ✓ | ✓ |
| beetEPRV2B-22 | 9 | 18612478 | 18608173 | 3510 | 794 | ✓ | n/a | n/a | ✓ | ✓ | ✓ |
| **beetEPRV2B-23\*** | **u1** | **201453** | **207019** | **4878** | **689** | ✓ | n/a | n/a | ✓ | ✓ | ✓ |
| **beetEPRV3-1\*** | **4** | **38153676** | **38161484** | **7513** | **296** | ✓ | ✓ | ✓ | ✓ | ✓ | ✓ |
| **beetEPRV3-2\*** | **5** | **15862348** | **15854689** | **7525** | **135** | ✓ | ✓ | ✓ | ✓ | ✓ | ✓ |
| **beetEPRV3-3\*** | **5** | **20484909** | **20492932** | **7513** | **511** | ✓ | ✓ | ✓ | ✓ | ✓ | ✓ |
| **beetEPRV3-4\*** | **5** | **55839215** | **55846739** | **n/a** | **0** | ✓ | ✓ | ✓ | ✓ | ✓ | ✓ |
| **beetEPRV3-5\*** | **7** | **45636054** | **45628533** | **7521** | **1** | ✓ | ✓ | ✓ | ✓ | ✓ | ✓ |
| beetEPRV3-6 | 7 | 47866227 | 47859680 | 6037 | 511 | ✓ | ✓ | ✓ | ✓ | ✓ | ✓ |
| **beetEPRV3-7\*** | **8** | **32243602** | **32236006** | **7522** | **75** | ✓ | ✓ | ✓ | ✓ | ✓ | ✓ |
| beetEPRV3-8 | 8 | 32257145 | 32250466 | 6680 | n/a | × | ✓ | ✓ | ✓ | ✓ | ✓ |
| **beetEPRV3-9\*** | **9** | **7568662** | **7561168** | **7495** | **0** | ✓ | ✓ | ✓ | ✓ | ✓ | ✓ |
| **beetEPRV3-10\*** | **u16** | **128580** | **120645** | **7479** | **457** | ✓ | ✓ | ✓ | ✓ | ✓ | ✓ |
| beetEPRV3-11 | u16 | 191123 | 184620 | 6504 | n/a | × | ✓ | ✓ | ✓ | ✓ | ✓ |

<sup>1</sup>The localization on an unplaced scaffold is marked by the prefix ‘u’. <sup>2</sup>Chromosomal positions given in reverse order indicate an inverse orientation of the respective sequence on the scaffold. <sup>3</sup>The beetEPRV length does not include the upstream terminal repeat (5’ TR). <sup>4</sup>The indication of an A or B next to the 5’ TR length implies a co-occurrence with the respective beetEPRV2 component.

We used the neighbor-joining algorithm (Saitou and Nei, 1987) embedded in *Geneious* 6.1.8 (https://www.geneious.com; Kearse *et al.*, 2012), for the reconstruction of the beetEPRV’s taxonomic affiliation to a known pararetroviral genus. Here, beetEPRV amino acid sequences were compared to the same EPRV reference set as for the *nhmmer* analysis. As outgroup the two sugar beet long terminal repeat (LTR) retrotransposons *Beetle*7 and Elbe2 of the Ty3-*gypsy* family were selected (Weber *et al.*, 2013; Wollrab *et al.*, 2012).

To assess the genomic environment of beetEPRVs copies individually, we visually inspected self dotplots of all members identified from sugar beet assembly and SMRT reads. Dotplots for a total of 514 sequences containing either MP or RT *nhmmer* hits or both of them with up to 8,000 bp flanking regions were generated automatically using the tool *FlexiDot* (Seibt *et al.*, 2018) with a wordsize of 9. With this method we were able to identify and analyze characteristic repetitive regions that appear up- and downstream of the full-length sequences, hereinafter called terminal repeats (TR).

We manually refined annotations of the beetEPRV members detected in the sugar beet assembly and the SMRT reads. Boxplots illustrating the sequence lengths of the beetEPRV members (derived from their chromosomal position; see Table 1) have been generated using ggplot2 (Wickham, 2009) implemented in R (R Core Team, 2018). The whiskers comprise all underlying data points.

### Search for beetEPRV transcripts and small RNA mapping

A publicly available cDNA library of *B. vulgaris* (genebank accession SRX674050) was searched for transcripts of the beetEPRV families using *blastn*. Small RNA reads (Zakrzewski *et al*., 2011) were mapped to the consensus sequences of the three beetEPRV families using the built-in mapping tool in *Geneious* 6.1.8 (https://www.geneious.com; Kearse *et al.*, 2012) with medium-low sensitivity and up to five iterations. Reads harboring insertions or deletions were discarded using a custom Python script. Read position, length, orientation, and counts were scored for graphical illustration by Python 2.7 using NumPy (Oliphant, 2006) and Matplotlib (Tosi, 2009).

### Plant material and genomic DNA extraction

Seeds of the *B. vulgaris* ssp. *vulgaris* genotype KWS 2320 were obtained from KWS Saat SE, Einbeck, Germany. Five other *Beta* and *Patellifolia* accessions as well as two further genera of the Amaranthaceae were analyzed: *B. vulgaris* ssp. *vulgaris* convar. *cicla* (chard ‘Vulkan’), *B. maritima* (BETA 1233), *B. patula* (BETA 548), *B. lomatogona* (BETA 674), and *B. nana* (BETA 541), *P. patellaris* (BETA 534), *Chenopodium quinoa* (CHEN 125), and *Spinacia oleracea* (‘Matador’). The respective seeds were obtained from the Leibniz Institute of Plant Genetics and Crop Plant Research Gatersleben, Germany. The plants were grown under long day conditions in a greenhouse. Genomic DNA was isolated from young leaves using the cetyltrimethylammonium bromide (CTAB) standard protocol (Saghai-Maroof *et al.*, 1984).

### PCR amplification, cloning and sequencing of beetEPRV3 sequences

Standard PCR reactions of genomic *B. vulgaris* DNA were performed using primer pairs designed for the RT and MP sequence of beetEPRV3 (see Table S2). The PCR conditions were 94 °C for 3 min, followed by 35 cycles of 94 °C for 1 min, primer specific annealing temperature for 30 sec, 72 °C for 45 sec, and a final incubation time at 72 °C for 5 min. PCR fragments were purified, cloned, and commercially sequenced. Sequenced inserts with an identity of at least 99.5 % to the reference beetEPRV3 element have been used as probes for the following hybridization experiments.

### Southern hybridization

Genomic DNA of sugar beet and related species was restricted with different enzymes, separated on 1.2 % agarose gels, and transferred onto membranes using alkaline transfer. We used random priming to radioactively label the beetEPRV3 probes (genebank accession LR812097-LR812098), followed by hybridization according to Sambrook *et al.* (1989). Filters were hybridized at 60 °C and washed at the same temperature in 2 × saline sodium citrate (SSC)/0.1 % sodium dodecyl sulfate (SDS) and 1 × SSC/0.1 % SDS for 10 min each. Signals were detected by autoradiography.

### Preparation of chromosome spreads

The meristem of young leaves was used for the preparation of mitotic chromosomes. For this, the leaves were treated for 3 h in 2 mM 8-hydroxyquinoline to accumulate metaphases, followed by fixation in 100 % methanol:glacial acetic acid (3:1). Fixed plant material was digested at 37 °C in the PINE enzyme mixture consisting of 2 % (w/v) cellulase from *Aspergillus niger* (Sigma C-1184), 4 % (w/v) cellulase ‘Onozuka R10’ (Sigma 16419), 2 % (w/v) cytohelicase from *Helix pomatia* (Serva C-8274), 0.5 % (w/v) pectolyase from *Aspergillus japonicus* (Sigma P3026), and 20 % (v/v) pectinase from *Aspergillus niger* (Sigma P4716) in citrate buffer (4 mM citric acid and 6 mM sodium citrate). After maceration, the mix was incubated for another 30 min and centrifuged at 4500 rpm for 5 min. The nuclei pellet was washed and resuspended in citrate buffer. To spread the chromosomes, 20 μl of the solution were dropped onto an ethanol-cleaned slide from a height of approximately 50 cm as published by Heslop-Harrison *et al.* (1991) and modified for beet by Schmidt *et al.* (1994). Finally, the chromosomes were rinsed in methanol:glacial acetic acid fixative.

### *Fluorescent* in situ *hybridization*

The beetEPRV3 probes (RT: genebank accession LR812097; MP: genebank accession LR812098) were labelled by PCR in the presence of digoxygenin-11-dUTP detected by antidigoxigenin-fluorescein isothiocyanate (FITC; both from Roche Diagnostics) and biotin-16-dUTP (Roche Diagnostics) detected by Streptavidin-Cy3 (Sigma-Aldrich), respectively. The probe pZR18S containing a part of the sugar beet 18S-5.8S-25S rRNA gene (HE578879; Dechyeva and Schmidt, 2009), as well as the probe pEV I marking an intercalary satellite DNA family (Schmidt *et al.*, 1991; Kubis *et al.*, 1998) were labelled with DY415-dUTP (Dyomics). The probe pXV1 for the 5S rRNA gene (Schmidt *et al.*, 1994) and the probe pBV I for the centromeric satellite DNA family (Schmidt and Metzlaff, 1991; Kubis *et al.*, 1998) were labelled with DY647-dUTP (Dyomics). Chromosomes were counterstained with DAPI (4’,6’-diamidino-2-phenylindole; Böhringer, Mannheim) and mounted in antifade solution (CitiFluor).

The hybridization and re-hybridization procedure was performed as described previously (Schmidt *et al.*, 1994) with a stringency of 82 %. Slides were examined with a fluorescence microscope (Zeiss Axioplan 2 imaging) equipped with appropriate filters. Images were acquired directly with the Applied Spectral Imaging v. 3.3 software coupled to the high-resolution CCD camera ASI BV300-20A. After separate capture for each fluorochrome, the individual images were combined computationally and processed using Adobe Photoshop CS5 software (Adobe Systems, San Jose, CA, USA). We used only contrast optimization, Gaussian and channel overlay functions affecting all pixels of the image equally. Chromosomes were identified and numbers assigned following Paesold *et al.* (2012). For this, we considered the position of the rRNA genes (pair one and four) and the distribution and density of the satellites pBV I and pEV I.

## RESULTS

### beetEPRVs can be grouped into three families according to their RT sequences

In order to identify endogenous pararetroviral sequences in the genome of sugar beet, we queried the high-quality *B. vulgaris* assembly EL10.1 (Funk *et al.*, 2018) for pararetroviral MPs and RTs (Hansen and Heslop-Harrison, 2004) using individual nucleotide Hidden Markov Models (nHMMs). As the RT is the key enzyme of all retroviral lineages (Xiong and Eickbush, 1990), we closely inspected all 119 RT matches. On nucleotide level, they showed identities of at least 50 % to the EPRV RT reference sequences (Llorens *et al.*, 2009; Llorens *et al.*, 2011; Becher *et al.*, 2014; Geering *et al.*, 2014) and were therefore assigned to the pararetroviruses. We also included truncated RTs with at least two of the seven conserved RT domains as defined by Xiong and Eickbush (1988) and Hansen and Heslop-Harrison (2004) in our dataset. To allow a sequence comparison, the pararetrovirus hits were aligned to each other; as outgroup, two Ty3-*gypsy* LTR retrotransposons from *B. vulgaris* were considered, *Beetle*7 and Elbe2 (Weber *et al.*, 2013; Wollrab *et al.*, 2012). A neighbor-joining tree (Fig. 1A) confirms a separation of all detected beet EPRV hits from the known Ty3-*gypsy* retrotransposons. The beetEPRV RTs form three distinct families marked by nucleotide sequence identities of less than 79 % between each other. In contrast, they can attain up to 100 % identity within one family, often reflected by short branch lengths (Fig. 1A, insets). We named the three families beetEPRV1, beetEPRV2, and beetEPRV3 according to the abundance of the pararetroviral sequences: The majority (50.4 %; n = 60) of the 119 beetEPRV RTs belongs to the family beetEPRV1, followed by beetEPRV2 with 30.3 % (n = 36) of the sequences. The smallest family is beetEPRV3 with 19.3 % (n = 23) of the assigned sequences.

**Fig. 1.**
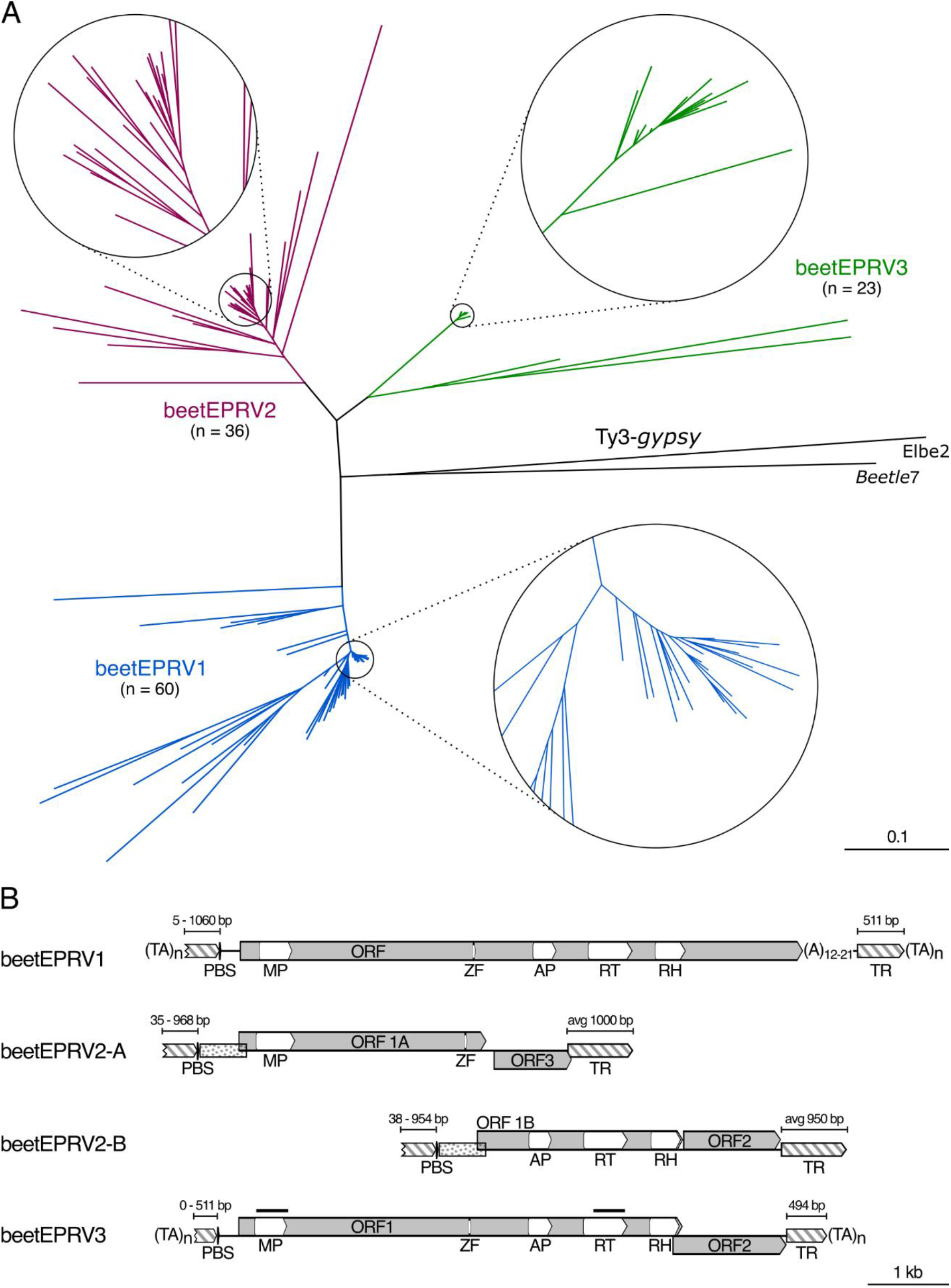
The diversity in sequence and structure of beet endogenous pararetroviruses (beetEPRVs) lead to their classification into three families. (A) Dendrogram showing the relationships among the 119 EPRV-RT hits found in the B. vulgaris genome. Two Ty3-gypsy RT sequences were used as outgroup (black), the chromovirus Beetle7 (genebank accession JX455085) and the errantivirus Elbe2 (genebank accession HE598759). Circles with zoomed dendrogram segments are shown for the densely packed branches. (B) Element structure of full-length beetEPRV consensus sequences from each family. Grey boxes mark open reading frames (ORFs); changes in the vertical position indicate frameshifts. White boxes within the ORFs represent conserved protein domains: MP = movement protein; ZF = zinc finger motif in the coat protein; AP = aspartic protease; RT = reverse transcriptase; and RH = RNase H1. The primer binding sites (PBS) complementary to the initiator tRNA of methionine (tRNA^Met^) are displayed. In addition, terminal repeats (TRs) are marked at the beginning and the end of the sequences as hatched boxes scaled according to their length. The conservation of the sequence between the PBS and the first ORF of both beetEPRV2 components is indicated as dotted boxes. As beetEPRV1 and beetEPRV3 sequences are usually terminated by TA microsatellites, the TA-rich sites are shown as (TA)_n_. Black bars above the MP and RT of the beetEPRV3 scheme indicate probes used for FISH.

To identify full-length beetEPRV members, we searched the RT hits for flanking upstream MP domains and further EPRV protein domains. In total, we detected 22 full-length sequences (see Table 1, asterisks), 14 beetEPRV1 and eight beetEPRV3 members with the EPRV-specific arrangement of the protein domains (MP-ZF-AP-RT-RH). Strikingly, no canonical beetEPRV2 sequences were found, neither in the genome assembly nor in the SMRT read data: Nearby beetEPRV2 MP domains (± 8 kb) were always separated from beetEPRV2 RT hits by additional primer binding sites (PBS; Supplementary data Fig. S2B).

The three beetEPRV families are characterized by distinct properties regarding their element structure, with beetEPRV1 and beetEPRV3 differing strongly from the structural organization of beetEPRV2 (Fig. 1B). Both beetEPRV1 and beetEPRV3 harbor all characteristic protein domains in an uninterrupted structure. BeetEPRV1 encodes a single, continuous ORF harboring all protein domains, whereas beetEPRV3 has two overlapping ORFs. The large beetEPRV1 ORF is terminated by a poly (A) region, while none of the identified beetEPRV3 ORFs has a poly (A) stretch at the 3’ end (see Fig. 1B). Both families contain a family-specific conserved region (approx. 500 bp) downstream of their ORF(s) that often is repeated in a fragmented manner upstream of the PBS (Fig. 1B, Supplementary data Fig. S2; hatched boxes). Due to its repeated nature and terminal position, we refer to it as terminal repeat (TR). All twenty beetEPRV1 members with an intact 5’ region (Table 1, column ‘PBS’) also contain at least five TR nucleotides upstream of the PBS, thus creating a conserved 5’-TATCC-3’ motif.

In contrast, beetEPRV2 members are organized differently: They exhibit a bipartite structure with the MP and ZF domain on one entity (component A, beetEPRV2-A) and the AP-RT-RH complex on the other (component B, beetEPRV2-B). As a consequence, most of the beetEPRV2 sequences including the AP-RT-RH polyprotein are located apart from the dedicated MP-ZF domain although co-occurrence of both entities has also been found (Supplementary data Fig. S2B). Both beetEPRV2 components also contain a non-functional ORF each (ORF3 and ORF2, respectively; Fig. 1B). Strikingly, both beetEPRV2 components are characterized by independent PBS motifs. Their underlying sequence 5’-TGGTATC(A/C)GAGC-3’ is homologous to the initiator tRNA methionine (tRNA^Met^) and the PBS of other EPRV genera (Hohn *et al.*, 1985; Verver *et al.*, 1987; Richert-Pöggeler and Shepherd, 1997). Similar to the poly(A) region of beetEPRV1, the 3’ TR of the beetEPRV2 components starts with an A-rich region that includes up to 31 adenines within a 34 bp window, potentially acting as polyadenylation signal.

### *All detected* B. vulgaris *EPRVs belong to the florendoviruses (FEVs)*

To assign the beetEPRV families to one of the characterized *Caulimoviridae* genera, we compared their consensus coding sequence of key domains (in particular RT and MP), to 13 selected EPRV representatives from eight plant EPRV genera, as well as the chromovirus *Beetle*7 and the errantivirus Elbe2 retrotransposons as outgroups (Fig. 2). We detected all seven conserved RT domains described by Xiong and Eickbush (1988) within each beetEPRV family (Fig. 2A): The average pairwise identity among the pararetroviral RTs is 46.9 %, within the genera it is above 60 % (CaMF and FMV; BSVAV and ComYMV; PVCV and FriEPRV), whereas to other genera and to the outgroup it is below 55 % and 38 %, respectively. Particularly, the beetEPRVs are most similar to the FEVs (69.2-79.3 %; Fig. 2A).

**Fig. 2.**
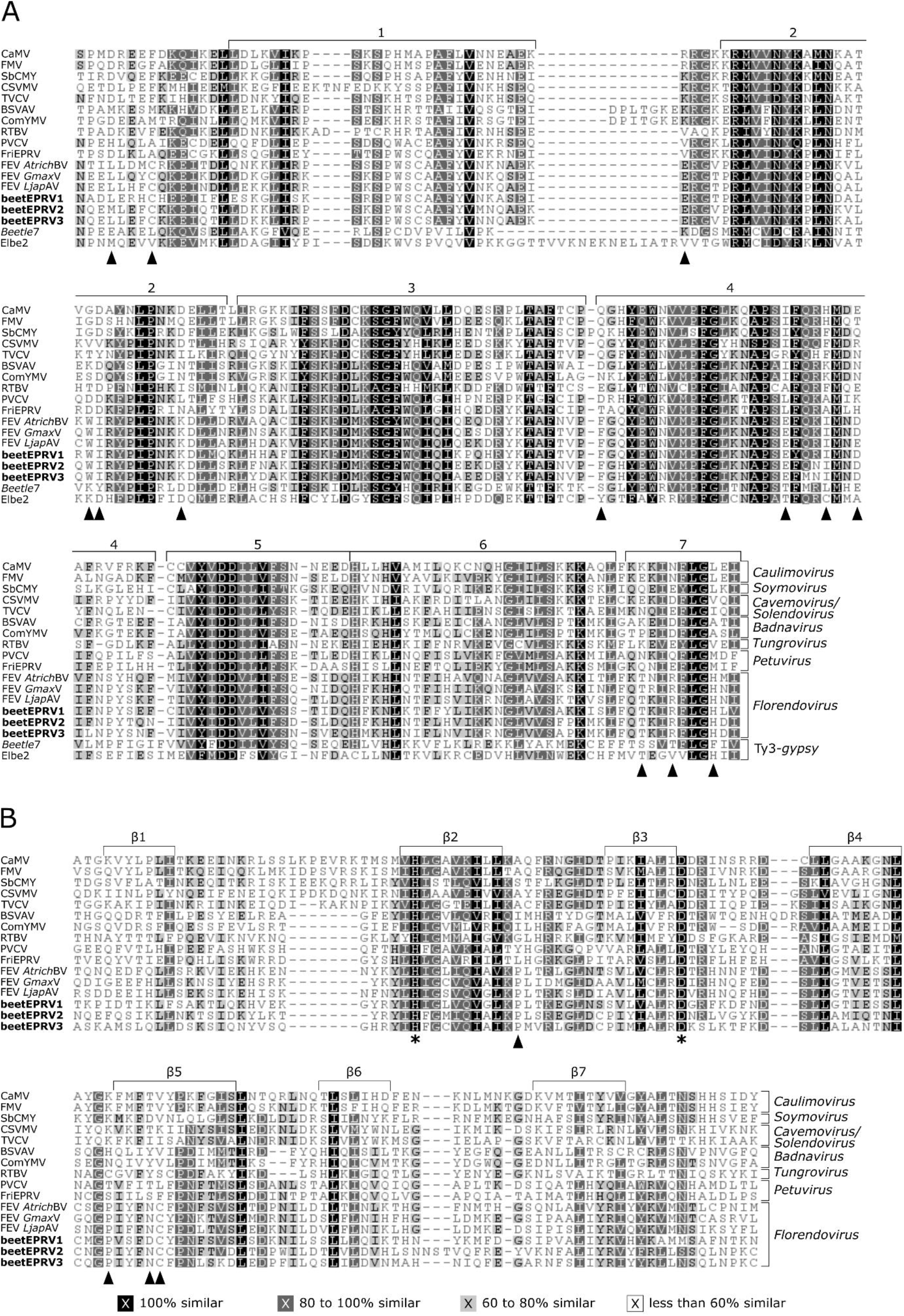
Comparative amino acid alignments of conserved EPRV protein domains compared to reference sequences. (A) EPRV and Ty3-gypsy RT and (B) EPRV MP sequence alignments accentuate high similarities between the beetEPRVs and the florendoviruses (FEVs). The shading reflects the similarity of the amino acids according to their physiochemical characteristics and FEV-characteristic amino acids are distinguished by triangles. (A) For the RT alignment, the Ty3-gypsy retrotransposons Beetle7 and Elbe2 were chosen as outgroup. Clamps above the sequences indicate the seven conserved RT domains. (B) Clamps above the sequences indicate the seven regions forming beta-strands in the MP secondary structure. Asterisks mark the conserved HX25D motif spanning the second and third beta-strand. Abbreviations of the reference elements: cauliflower mosaic virus (CaMV) and figwort mosaic virus (FMV) from the genus Caulimovirus, soybean chlorotic mottle virus (SbCMV; Soymovirus), cassava vein mosaic virus (CSVMV; Cavemovirus) and tobacco vein clearing virus (TVCV; Solendovirus), banana streak VA virus (BSVAV) and Commelina yellow mottle virus (ComYMV) from the genus Badnavirus, rice tungro bacilliform virus (RTBV; Tungrovirus), petunia vein clearing virus (PVCV) and Fritillaria imperialis EPRV (FriEPRV) from the genus Petuvirus, and the florendoviruses (FEV) Amborella trichopoda B virus (AtrichBV), Glycine max virus (GmaxV), and Lotus japonicus A virus (LjapAV)

The MP core is characterized by seven beta-strands (Fig. 2B; Mushegian and Elena, 2015) which together form a secondary structure similar in all viral MPs (validated by the internet tool JPred 4.0; Drozdetskiy *et al.*, 2015). We found a conserved HX_25_D motif that extends to the second and third beta-strand of all analyzed EPRV MPs (Fig. 2B, asterisks). Remarkably, an insertion of three amino acids between the sixth and seventh beta-strand of the MP (Fig. 2B) is unique for beetEPRV2 and sets this family apart from beetEPRV1 and beetEPRV3. In general, the MP is more variable than the other pararetroviral protein domains (ZF, AP, RT, RH), and the average pairwise nucleotide identity amounts to just 25.4 %; within the genera we found a minimum of 36 % and between genera a maximum of 36 %. Again, the beetEPRV MPs show the highest identity to the representative FEVs with a value of 36-50 %. In both alignments, we detected discriminatory amino acids present in all analyzed FEVs including the beetEPRVs, which distinguished them from the other EPRV genera (Fig. 2; triangles; Table S3). Differences in the number of amino acids separating two domains from another can also be used to distinguish genera from each other.

Neighbor-joining dendrograms based on the RT and MP alignments are similar. The affiliation of the beetEPRV families to the FEVs is demonstrated by a maximum bootstrap support of 100 % (Fig. 3). However, despite their common host, beetEPRV1 is not grouped on the same branch with the other two beetEPRV families. In the RT analysis, beetEPRV1 is in a basal position relative to the other FEVs, including the FEV *Atrich*BV whose host is the evolutionarily old, basal angiosperm *Amborella trichopoda*. In contrast, in the MP analysis this basal position is taken by beetEPRV2 and beetEPRV3, indicating a high structural diversity in the analyzed beetEPRVs.

**Fig. 3.**
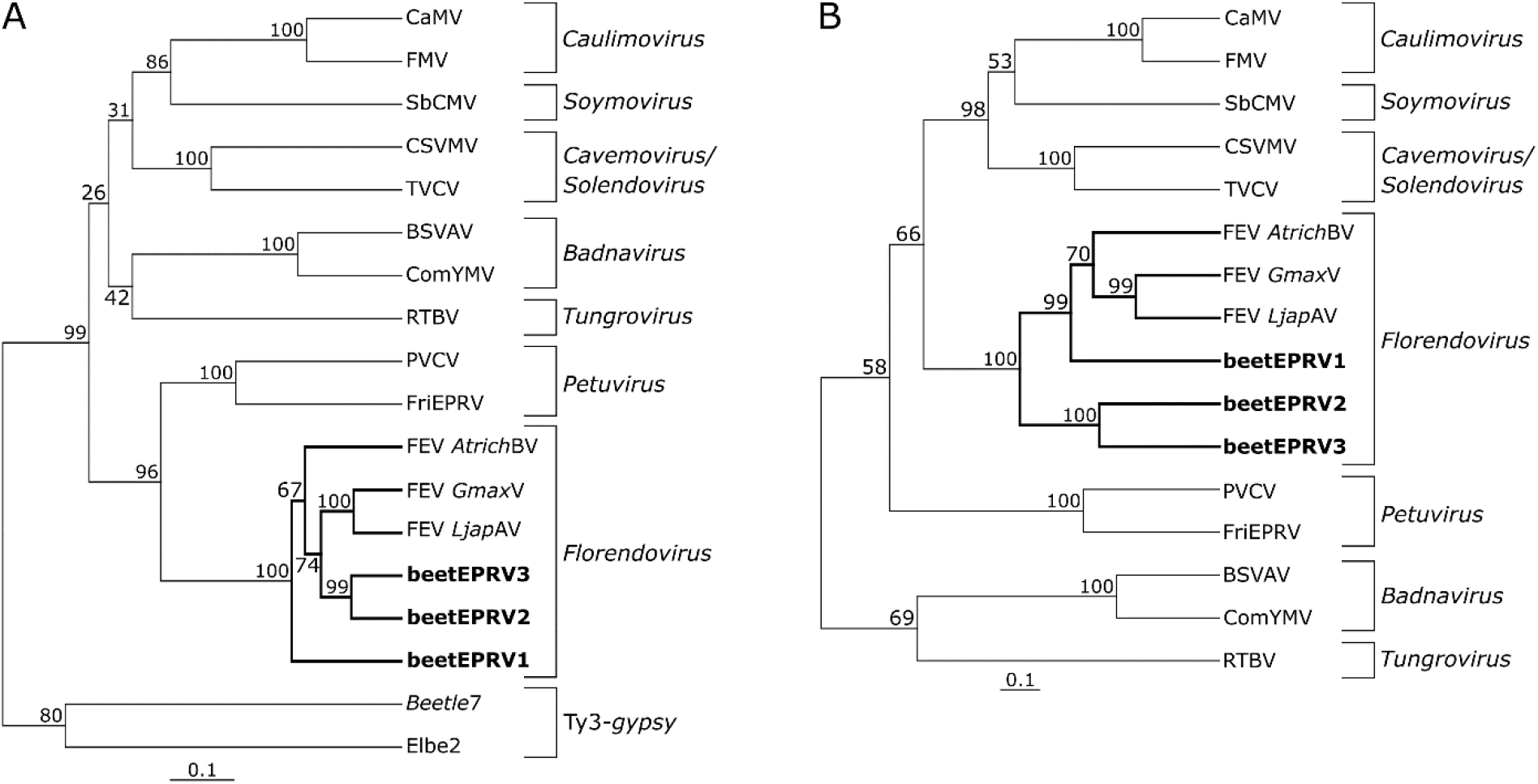
Dendrograms grouping the beetEPRVs to the florendoviruses based on the protein sequence of reverse transcriptase, RT (A) and movement protein, MP (B). Dendrograms were constructed with the neighbor-joining method. Bootstrap values (1,000 replicates) are shown above the nodes. This analysis includes the RT and MP domains of the representative beetEPRV sequences (see Fig. 1B) and the homologous region of the caulimoviruses and Ty3-gypsy retrotransposons as references (for abbreviations see legend of Fig. 2).

Apart from the sequence similarities of the RT and MP, a number of further beetEPRV hallmarks support their assignment to the FEVs: With 7.6 kb and 7.5 kb, respectively (see Table 1; Fig. 4A), the mean length of beetEPRV1 and beetEPRV3 members corresponds to the length of FEVs (7.2-8.5 kb; Geering *et al.*, 2014). Due to an additional ORF3, composite beetEPRV2 elements were much longer (10.7 kb; component A = 5.5 kb and component B = 5.2 kb). The additional ORF2 in beetEPRV2-B and beetEPRV3, is also characteristic for FEVs and is assumed to encode an FEV-specific protein. Its estimated molecular weight of 50-54 kDa is well in line with the ORF2 of other FEVs (45-58 kDa; Geering *et al.*, 2014). Although there is no clear ORF2 in the beetEPRV1 reference element due to the accumulation of frameshifts, there are several AUG start codons within a short interval in the 3’ region of ORF1 that enable reconstruction of a putative ORF of an appropriate molecular weight (51-55 kDa, depending on the start codon position).

**Fig. 4.**
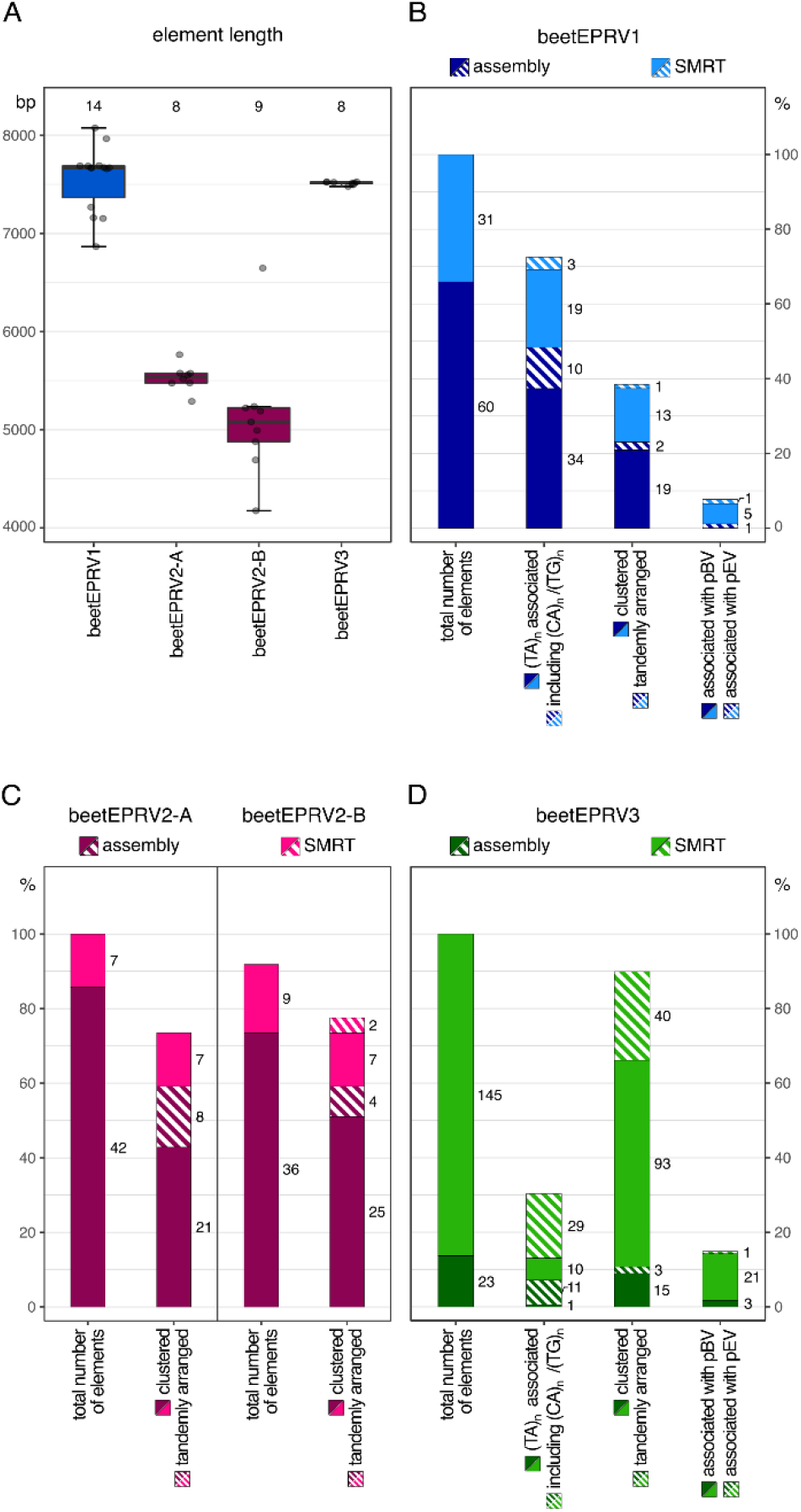
Structural characteristics of the three beetEPRV families including mean lengths (A) and genomic contexts (B-D). (A) Mean length of beetEPRV1, the two components beetEPRV2-A and beetEPRV2-B, and beetEPRV3. The underlying sequences refer to elements marked by an asterisk in Table 1. (B-D) Analysis of the genomic context of beetEPRV1 (B), beetEPRV2 (C), and beetEPRV3 (D) using two sequence data sets of the sugar beet genotype EL10 (assembly: dark color; SMRT reads: light color). As beetEPRV1 (B) and beetEPRV3 (D) showed a greater variation in properties compared to beetEPRV2-A (C left) and beetEPRV2-B (C right), the corresponding barcharts highlight different aspects: (B, D) The first bar represents the total beetEPRV number (100 %). The second bar shows an association with various micro-satellites. The third bar demonstrates the amount of clustered/nested (filled) and tandemly arranged (striped) beetEPRV sequences. The fourth bar shows an association to known beet tandem repeats, namely the centromeric pBV (filled) as well as the intercalary pEV (striped) satellite repeat. (C) The first bar represents the total beetEPRV2-A number (100 %). The second bar demonstrates the amount of clustered/nested (filled) and tandemly arranged (striped) beetEPRV2-A sequences. The third bar represents the total beetEPRV2-B number (100 %). The fourth bar demonstrates the amount of clustered/nested (filled) and tandemly arranged (striped) beetEPRV2-B sequences. The sample size is given above the boxplots (A) and next to the barcharts (B-D). The percentages (B-D) refer to the total number of analyzed beetEPRV sequences of the respective family.

Taken together, based on the sequence similarities in the key protein domains, the conserved element length, and the presence of an additional ORF, we conclude with confidence that all detected EPRVs in beet belong to the FEV genus.

### beetEPRVs are embedded in a repeat-rich environment

To assess the genomic context, we manually extracted 514 individual candidate beetEPRV sequences (full-length as well as partial) from the sugar beet genotype EL10 reference genome assembly and the corresponding SMRT read data (Funk *et al.*, 2018). Using our nHMMs, we extracted:

- 161 sequences from the EL10 assembly with 60 beetEPRV1, 42 beetEPRV2-A, 36 beetEPRV2-B, and 23 beetEPRV3 sequences
- 353 sequences from the raw EL10 PacBio long (SMRT) reads with 31 beetEPRV1, 7 beetEPRV2-A, 9 beetEPRV2-B, and 306 beetEPRV3 sequences.

Differences in the relative abundance of the three beetEPRV families, such as the high abundance of beetEPRV3 in the long reads but its rareness in the assembled genome, may reflect biases in these two datasets.

We compared these pararetroviral elements to the respective consensus sequences (Fig. 1B; Supplementary data S1), examined self dotplots to investigate each element’s structure and organization, annotated the beginning and the end of each integrated pararetrovirus (Fig. 4A) and investigated the flanking regions (about 8 kb each site). The majority (73 % from the assembly; 71 % from the SMRT reads) of the beetEPRV1 sequences as well as several beetEPRV3 sequences (52 % and 27 %, respectively) are directly flanked by AT-rich low complexity regions at one or both ends. These are often arranged as (TA)n microsatellites (Fig. 4B and 4D) that frequently harbor short stretches of CA or TG dinucleotides. In some cases, beetEPRV3 elements on the SMRT reads are flanked by longer motifs such as TATC (n = 1), TATACA (n = 5) and TTTCGGGG (n = 1). In contrast, we did not detect any beetEPRV2 members associated with low complexity motifs.

BeetEPRV1, beetEPRV2, and beetEPRV3 members without low complexity (TA)n flanking regions were also detected in highly repetitive neighborhood, characterized by fragmental duplications, rearrangements and juxtapositions of further truncated beetEPRV copies. In particular, beetEPRV elements of the same family often localize close to each other as fragments or full-length copies. Noteworthy, the intact, adjacent beetEPRV sequences are connected by TRs (Supplementary data Fig. S2). Taking into account the assembly and the SMRT reads, the frequency of such arrangements in tandem-like arrays varied with 3 % for beetEPRV1, 19 % for beetEPRV2-A, 11-22 % for beetEPRV2-B, and 13-28 % for beetEPRV3 (Fig. 4B-D). Regarding the bipartite nature of beetEPRV2, a tandem-like arrangement of the beetEPRV2 components A and B in the same orientation was found frequently (assembly: 33 %, SMRT: 63 %), in which the A-B arrangement was half as common as the B-A arrangement (Supplementary data Fig. S2).

Some beetEPRV1 sequences (2-3 %) localize adjacent to units of the intercalary satellite DNA family pEV I described by Schmidt *et al.* (1991), while more (16 %; Fig. 4B) border the centromeric pBV satellite arrays of Schmidt and Metzlaff (1991). BeetEPRV3 was also found to be associated with pEV I and pBV (0.7 % and 13-14 %, respectively; Fig. 4D). For two instances along the assembly and five instances on the SMRT reads, beetEPRV3 was flanked by pBV arrays on both sides. In addition, combinations of a pBV array on one end and a (TA)_n_ microsatellite on the other were also detected. All six described pBV subfamilies (Zakrzewski *et al.*, 2013) were observed in the associations with beetEPRV1 and beetEPRV3.

In summary, the sugar beet EPRVs are embedded in highly repetitive genomic contexts. They form complex clusters, often containing multiple, rearranged elements of the same beetEPRV family. Co-occurrences with heterochromatic satellite DNAs as well as low complexity microsatellites were observed frequently.

### beetEPRVs show a high, locally focused coverage with small RNAs

It is assumed that most repeats are transcribed at a basal level regulated by the host through epigenetic silencing (reviewed by Lippman and Martienssen, 2004). We detected transcripts for all three beetEPRV families in a cDNA library (genebank accession SRX674050) that could potentially lead to virus activation. To investigate how the beet host genome may prevent such virus activation, we analyzed the potential silencing by small RNAs (smRNAs). Publicly available smRNA reads from *B. vulgaris* were mapped against the consensus sequences of the three beetEPRV families (see Fig. 1B; Supplementary data S1). Out of 20,091,021 smRNA reads in total, 1,051 reads matched to beetEPRV1, 381 reads to the concatenated consensus sequence of both beetEPRV2 components, and 13,235 reads to beetEPRV3 (Fig. 5). Among the smRNAs matching the three beetEPRV families, only beetEPRV3-derived smRNAs comprised the wide spectrum from 18 to 30 nt (Fig. 5A). However, for all three beetEPRV families, smRNAs with a length of 20 to 26 nt contributed more than 99 % of all mapping smRNAs. These smRNAs were divided in smRNAs that induce posttranscriptional gene silencing (PTGS, 20-23 nt; Rosa *et al.*, 2018) and transcriptional gene silencing (TGS, 24-26 nt; Ghoshal and Sanfaçon, 2015). In beetEPRV1 and beetEPRV3, about two third of the smRNAs potentially mediates PTGS (65.9 % and 64.7 % respectively), whereas TGS-associated smRNAs contribute the smaller fraction (33.7 % and 34.8 % respectively). Strikingly, the opposite applies for beetEPRV2: Only 13.4 % of the smRNAs potentially induce PTGS, while 85.8 % may lead to TGS.

**Fig. 5.**
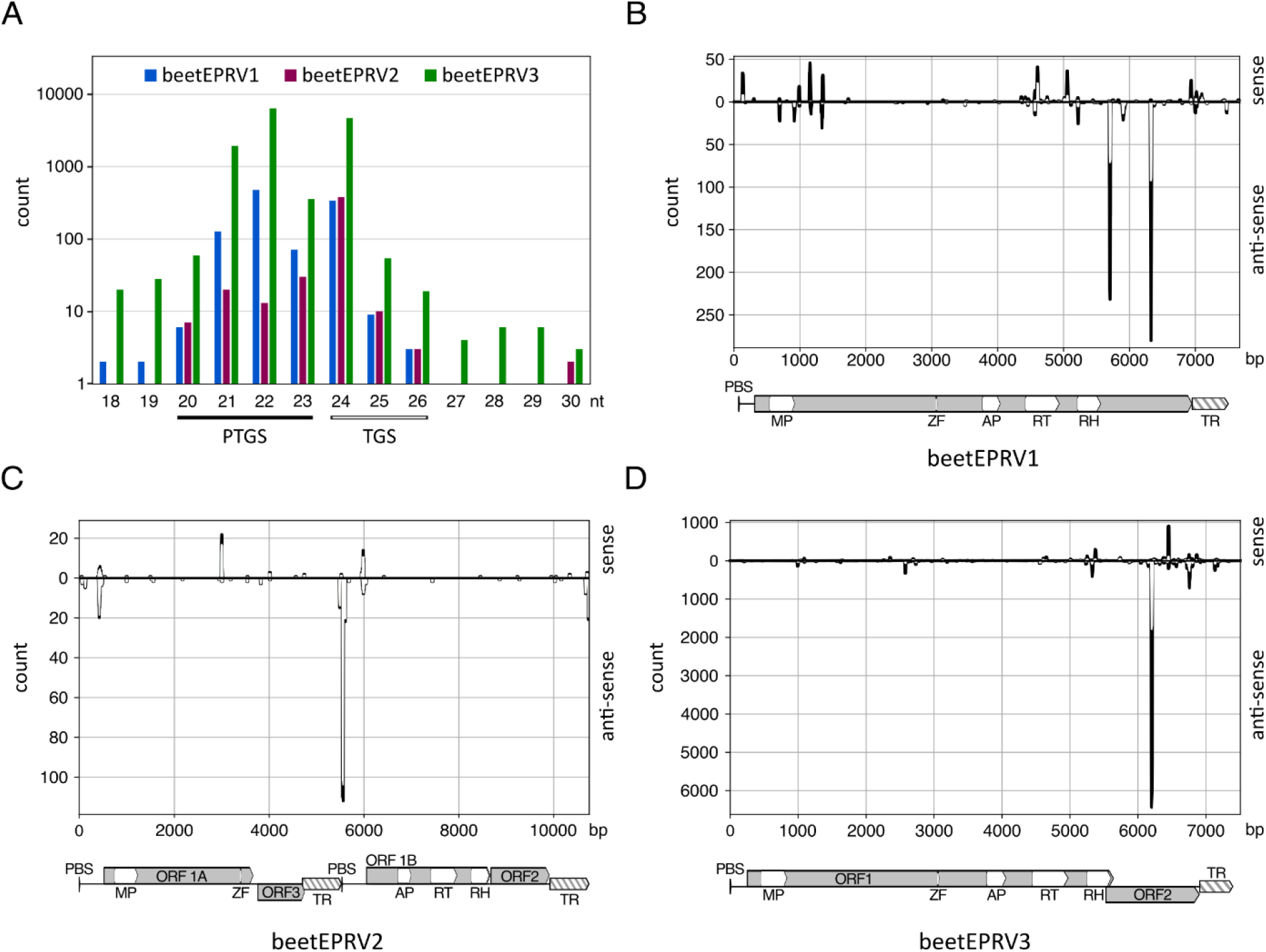
Evaluation of small RNAs (smRNAs) mapping to the consensus sequences of the three beetEPRV families. Small RNAs inducing posttranscriptional gene silencing (PTGS; 20-23 nt) are marked by black bars and those inducing transcriptional gene silencing (TGS; 24-26 nt) by white bars. (A) Length distribution of smRNAs matching the respective beetEPRV consensus sequences (color-coded bars). Note the logarithmic scale. (B-D) Read depths of smRNAs mapped against the beetEPRV consensus sequences. Positive peaks represent sense RNAs, negative peaks indicate anti-sense RNAs. Only smRNAs mediating PTGS (black; 20-23 nt) and TGS (white; 24-26 nt) have been considered. The respective beetEPRV element structure is shown below the diagram as introduced in Fig. 1B.

We observed peaks with high (>100 reads, beetEPRV2) and very high smRNA read abundances (>200 reads, beetEPRV1; >6,000 reads, beetEPRV3) at particular positions along the three consensus sequences (Fig. 5B-D). These peaks are preferentially located in regions or ORFs that do not carry any known protein domains: This refers to ORF2 in beetEPRV3 and the 3’ region of the beetEPRV1-ORF, as well as to beetEPRV2’s internal region between ORF1A and ORF1B. Moderately sized peaks were also found in the terminal repeats.

### *beetEPRV3 is highly methylated in the* B. vulgaris *genome and occurs in closely related beet species*

As beetEPRV3 is characterized by the highest average pairwise identity of its members, many intact copies with continuous ORFs, and as we found an extraordinarily high copy number in the SMRT read data set, we used this family as reference for experimental studies on EPRVs in beet. In order to gain information about cytosine methylation of beetEPRV3 sequences, we restricted genomic *B. vulgaris* DNA with the methylation-sensitive enzymes *Msp*I and *Hpa*II and hybridized the RT and MP probe to the membranes (Fig. 6A and 6B, lanes 6 and 7; Table S2). Whereas *Hpa*II only cuts unmethylated CCGG sequences, *Msp*I is able to tolerate methylation of the internal cytosine (Waalwijk and Flavell, 1978). In the beetEPRV3 reference sequence, the CCGG restriction site is present three times; point mutations may also lead to further or fewer CCGG sites. As beetEPRV3 was neither cut by *Hpa*II nor *Msp*I, we conclude methylation of the outer or both cytosines occurred within the CCGG motifs.

**Fig. 6.**
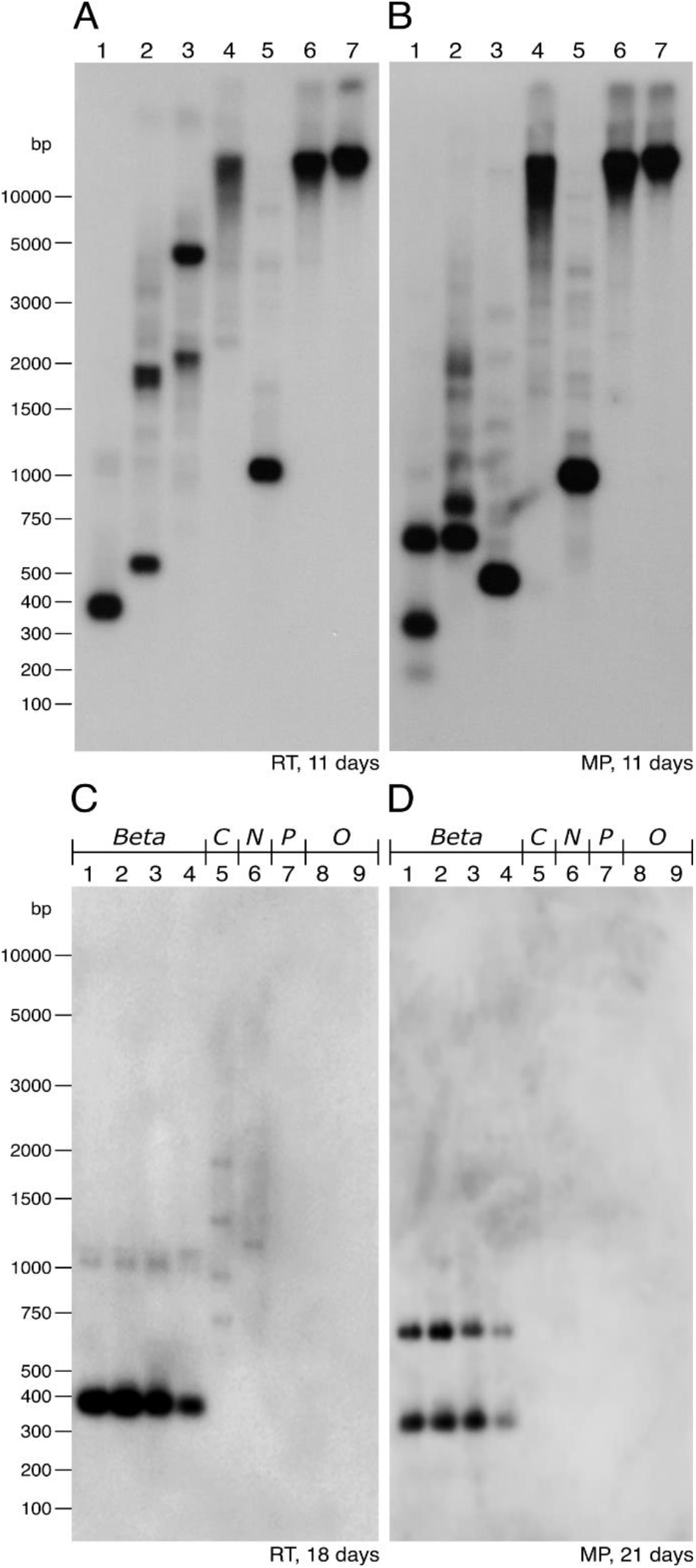
Autoradiograms of the Southern hybridization to estimate the abundance of beetEPRV3 in sugar beet and related species. As probes reverse transcriptase RT (A, C) and movement protein MP (B, D) were used. For (A, B), DNA of B. vulgaris was restricted by different enzymes: AluI (1), FspBI (2), BseGI (3), BspCNI (4), HindIII (5), MspI (6) and HpaII (7). For (C, D), AluI restricted DNA of different Amaranthaceae species was used: From the section Beta the B. vulgaris cultivars KWS 2320 (1) and Swiss chard (2), B. maritima (3), and B. patula (4); from the section Corollinae (C) B. lomatogona (5); from the section Nanae (N) B. nana (6); and from the sister genus Patellifolia patellaris (P, 7); quinoa (Chenopodium quinoa; 8), and spinach (Spinacia oleracea; 9) were selected as outgroup (O). Hybridization was carried out with a stringency of 79 %.

The genomic *B. vulgaris* DNA (KWS 2320) was also restricted by five additional restriction enzymes to estimate the sequence conservation and abundance of beetEPRV3 in sugar beet (Fig. 6A and 6B; lanes 1-5). The Southern hybridization required a long exposition time (11 days), a sign for a low beetEPRV3 abundance in *B. vulgaris*. The clear bands point to a strong conservation of the restriction sites within the beetEPRV3 sequence, confirming the high similarity of beetEPRV3 members to each other.

In order to investigate the beetEPRV3 abundance in related genomes, we comparatively hybridized both probes (RT and MP) to *Alu*I-restricted DNA of the *B. vulgaris* cultivars KWS 2320 and Swiss chard, *B. maritima*, *B. patula*, *B. lomatogona*, *B. nana*, as well as *P. patellaris*, a member of the *Beta* sister genus *Patellifolia* (Fig. 6C and 6D). As outgroup we chose *Chenopodium quinoa* and *Spinacia oleracea*, both also belonging to the Amaranthaceae. The two probes hybridized to all four genomes of the section *Beta*, producing the expected *Alu*I patterns (Fig. 6C and 6D, lanes 1-4). This included the cultivated beet and chard species as well as the wild beet *B. maritima*. In the further sections of the genus Beta represented by *B. lomatogona* and *B. nana*, RT hybridization signals also became visible, but in a divergent pattern with lower intensity. Signals for the MP probe, which were less conserved than the RT in EPRVs (see Fig. 2) were not observed, even after an extended exposure time. This may indicate either beetEPRV3 presence in much lower abundance and/or higher divergence in these species or, more likely, cross-hybridization of the RT probe to a related pararetroviral sequence. In the sister genus *Patellifolia* as well as in the outgroups no signals were detected, neither for the RT nor for the MP probe. This indicates no significant sequence homologies between beetEPRV3 from *B. vulgaris* and possible EPRVs from *P. patellaris*, quinoa, and spinach.

### *beetEPRV3 clusters on all* B. vulgaris *chromosomes*

To determine the chromosomal localization of beetEPRV3, mitotic chromosomes of *B. vulgaris* (KWS 2320) were prepared and hybridized with the biotin-labelled beetEPRV3 RT probes (red) and digoxygenin-labelled beetEPRV3 MP probes (green; Fig. 1B; Fig. 7A, B; Supplementary data Fig. S3). The RT and MP signals for beetEPRV3 often co-localize, shown by the merging of red and green signals to yellow ones in the overlay (Fig. 7D). Nevertheless, the observation of distinct signals may point to the presence of truncated beetEPRV3 sequences. This observation is corroborated by the bioinformatic analyses in which the reshuffling of beetEPRV sequences was also detected.

**Fig. 7.**
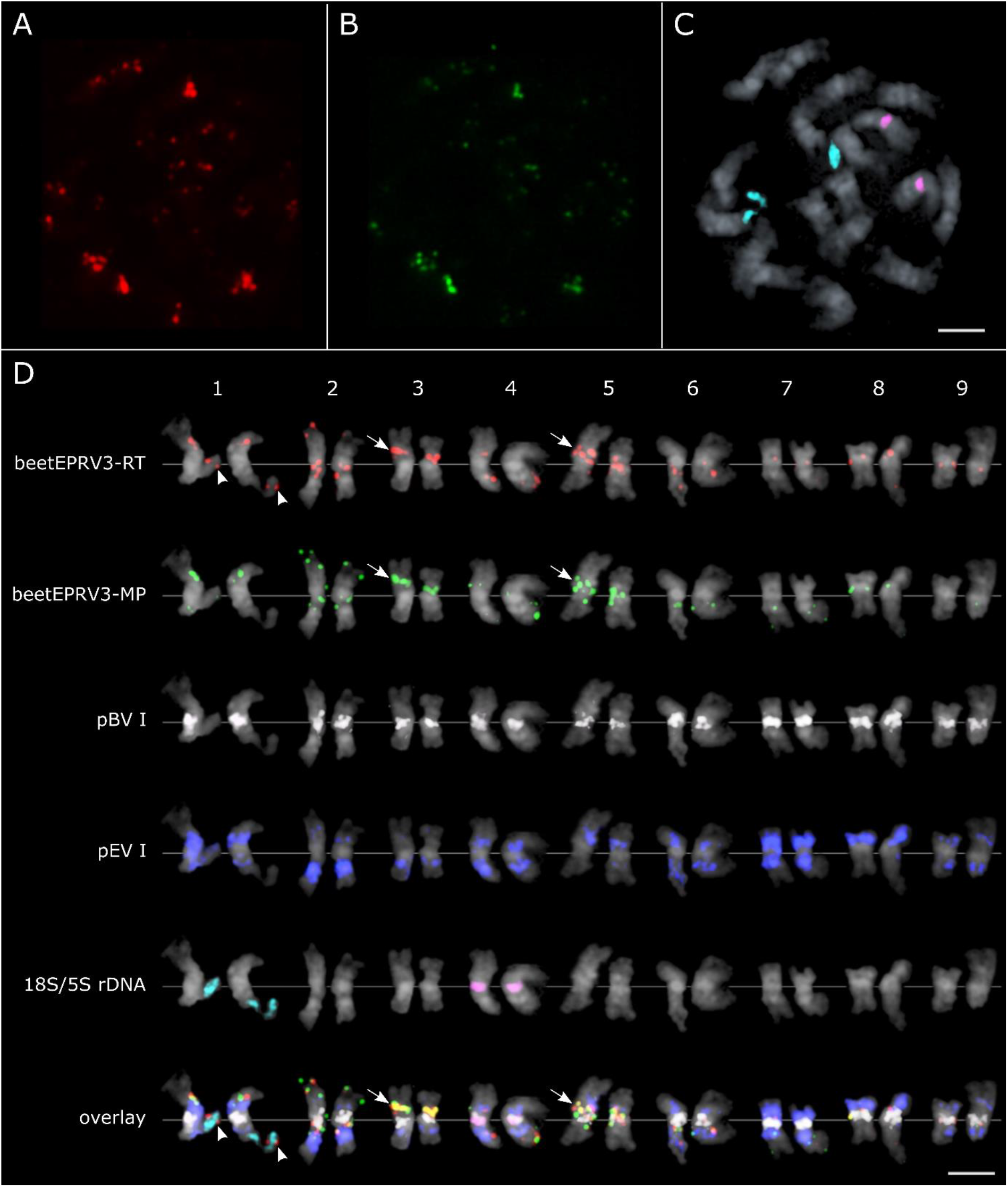
Localization of beetEPRV3 along mitotic metaphase chromosomes of B. vulgaris. DAPI-stained mitotic chromosomes of B. vulgaris are shown in grey. Cloned sequences of the RT (red) and the MP (green) domain of beetEPRV3 were used as probes. (A-D) Multicolour FISH of beetEPRV3-RT (red), beetEPRV3-MP (green), centromeric pBV I (white), intercalary pEV I satellite (blue), 18S rDNA genes (turquoise), and 5S rDNA genes (magenta). Information on probe labelling and detection can be found in the materials and methods section. (D) Sorted chromosomes from Fig. 6 A-C additionally showing pBV and pEV signals. Paired chromosomes represent the homologous chromosomes. The assignment of chromosome numbers is based on the rDNA genes (pair one and four) and on the distribution and density of the satellites pBV I and pEV I according to Paesold et al., 2012. The strongest beetEPRV3 clusters on chromosomes 3 and 5 are highlighted (arrows), as well as the co-localization of beetEPRV3 fragments, including the RT, with the 18S-5.8S-25S rDNA (arrowheads). Bars = 2 μm

The diploid chromosome set of *B. vulgaris* consists of 18 chromosomes and all chromosomes can be identified using rDNA and repetitive sequences (Schmidt *et al.*, 1994; Paesold *et al.*, 2012). Therefore, the beetEPRV probes were hybridized together with additional probes for the 18S-5.8S-25S and 5S rDNA (turquoise and magenta in Fig. 7C, 7D and Supplementary data Fig. S3) to identify the homologous chromosomes 1 and 4, respectively. To designate the remaining chromosomes, a re-hybridization with probes marking the centromeric satellite DNA pBV I and the intercalary satellite DNA pEV I (white and blue in Fig. 7D and Supplementary data Fig. S3) was carried out. The intensity and co-occurrence of these satellite arrays allowed the assignment to the respective chromosome pairs according to Paesold *et al.*, 2012.

Hybridization of beetEPRV3 RT and beetEPRV3 MP probes show the localization of beetEPRV3 on all 18 *B. vulgaris* chromosomes (Fig. 7D; Supplementary data Fig. S3). Signal strengths differ strongly between chromosomes, ranging from very strong to faint fluorescence, indicating the presence of large and small accumulations of beetEPRV3 sequences and the tendency to form clusters. The signals were often detected in heterochromatic regions which is revealed by the co-localization with the heterochromatic satellite DNAs (pBV I, pEV I) and densely stained DAPI signals (Supplementary data Fig. S3). This is in accordance with the association with these sequences found during the analysis of the flanking sequences. We detected pericentromeric, intercalary and distal positions with similar localization patterns along the two homologous chromosomes. Thus, the strongest clusters reside in the intercalary region of chromosome 3 and the pericentromeric region of chromosome 5 (Fig. 7D, arrows). FISH images of the chromosomes 1 and 2 show the most distal signals, where we detected a co-localization of beetEPRV3 RT with the 18S-5.8S-25S rDNA (Fig. 7D, arrowheads).

## DISCUSSION

EPRVs are a widespread component of plant genomes that often accumulate over time and become a noteworthy part of the repetitive fraction of the genome (Hohn *et al.*, 2008; Diop *et al.*, 2018; Gong and Han, 2018). Our analyses in sugar beet show that EPRVs make up approx. 0.3 % of the *B. vulgaris* genome, a value being within the EPRV proportion range of 0-2 % in other host plants (Geering *et al.*, 2014; Duroy *et al.*, 2016). Although we find some essentially intact representative EPRV sequences, the majority of the pararetroviral sequences in beet do not comprise all EPRV-specific protein domains due to fragmentation and/or truncation. For this reason, it is likely that some beetEPRV fragments with strongly diverged or missing RT or MP protein domains (the major criteria used to search for EPRVs and applied here) may have escaped our computational detection and that the aforementioned beetEPRV quantification is probably an underrepresentation. Nevertheless, the resulting, relatively large amount of integrated EPRV sequences was unexpected given that no exogenous viral sequences have been reported for beets so far.

Based on their sequence and structural characteristics, we subdivided the sugar beet EPRVs into three families: beetEPRV1, beetEPRV2, and beetEPRV3 (Fig. 1). Considering their overall length, their structure with a putative additional ORF, and their amino acid sequence homologies, all beetEPRV families represent typical members of the genus *Florendovirus* (FEV) as described by Geering *et al.* (2014).

### The family-specific element structure indicates different evolutionary beetEPRV origins

Certain features in beetEPRV structure and sequence provide evidence for potential infections and subsequent amplifications at multiple time points during beet evolution: For the reference family beetEPRV3, we detected high nucleotide identities between the individual beetEPRV3 members and multiple genomic copies with intact open reading frames. Comparative Southern hybridizations with MP and RT probes led to conserved patterns within the section *Beta*, indicative of a single or only few integration events that were subsequently amplified within the beet genome.

The species in the sister sections *Corollinae* and *Nanae* do not seem to harbor beetEPRV3-related sequences as they do not produce MP signals and only weak, dissimilar signals for the RT, likely the result of cross-hybridization rather than true homology. Together, this may point to an initial beetEPRV3 integration into the beet genome after the split of the sections *Corollinae/Nanae* from *Beta* approx. 13.4 to 7.2 million years ago (mya; Hohmann *et al.*, 2006). Given the estimated age of FEVs (34-20 mya; Geering *et al.*, 2014) and EPRVs in general (320 mya; Diop *et al.*, 2018) this seems to be an evolutionarily young infection history. Yet, in comparison with the time of invasion of EPRVs into other host species (e.g. eBSV into the genome of *Musa* sp. 640,000 years ago, Gayral *et al.*, 2010, Duroy *et al.*, 2016; eRTBVL-D into the genome of *Oryza* sp. 2.4-15 mya, Chen *et al.*, 2018) or compared with the estimated integration time points of other retroelements into the sugar beet genome (chromovirus *Beetle*2: 130,000 years ago, Weber and Schmidt, 2009; Cassandra TRIMs: 0.1-8 mya, Maiwald *et al.*, 2020) the assumed beetEPRV3 endogenization event ranges at a similar timeline. From its most basal position within the FEVs in the RT dendrogram (Fig. 3A), we believe that the beetEPRV1 family may comprise the oldest beetEPRV members. Its element structure is also less complex compared to the classical FEV elements: Although FEVs typically have two overlapping ORFs (Geering *et al.*, 2014), beetEPRV1 contains a single, continuous ORF that may be a sign of an early stage in FEV evolution. Nevertheless, despite the low complexity of beetEPRV1, we do not think that this family served as precursor for beetEPRV2 and beetEPRV3. Instead, as the beetEPRV1 ORF differs considerably from those of the other beetEPRV families and all dendrograms place beetEPRV1 separately (Fig. 1 and 3), we argue for an independent beetEPRV1 infection event.

In contrast to beetEPRV1 and beetEPRV3, elements of the beetEPRV2 family are characterized by a bipartite structure with the two components beetEPRV2-A and beetEPRV2-B. The components have the same 5’ end and their first ORFs exhibit a high overall similarity to the respective regions of beetEPRV1 and beetEPRV3 (Fig 1B and Fig. 8A, dotted boxes). This supports an origin from a beetEPRV2 precursor containing both A and B components in a single element (Fig. 8B).

**Fig. 8.**
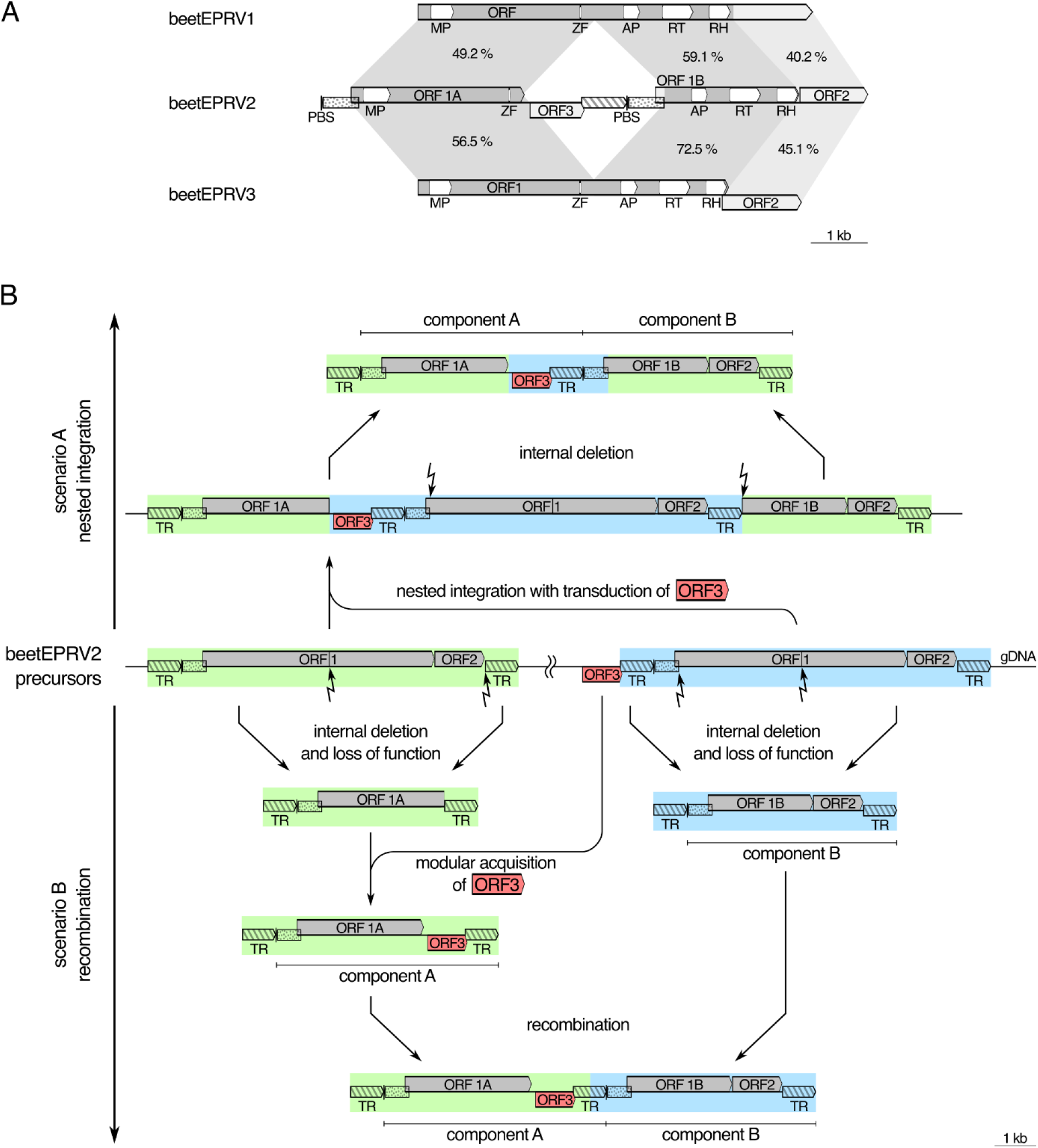
Evolutionary hypothesis to explain the bipartite structure of beetEPRV2 based on sequence similarities to the other beetEPRV families. (A) Similarity of the coding regions of beetEPRV2 to beetEPRV1 and beetEPRV3. The nucleic acid sequence identity between the ORF sections is given in the respective grey-shaded regions. BeetEPRV2’s internal region between ORF1A and ORF1B does not have an equivalent in beetEPRV1 and beetEPRV3. (B) We propose two evolutionary scenarios that may have led to the bipartite structure of beetEPRV2: In scenario A, we assume the nested integration of two beetEPRV2 precursor sequences. The internal beetEPRV2 element may have piggybacked an additional ORF (ORF3). To yield the bipartite element observed today, internal domains were subsequently deleted. Scenario B may have involved the recombination of two independently reshuffled and partly deleted beetEPRV2 entities. ORF = open reading frame; MP = movement protein; ZF = zinc finger in the coat protein; AP = aspartic protease; RT = reverse transcriptase; RH = RNase H1; PBS = primer binding site; TR = terminal repeat; dotted boxes = homologous sequence regions.

There are different possible scenarios to explain the bipartite nature of beetEPRV2 and its duplicated 5’-end (Fig. 8B): On the one hand, nested integration of one beetEPRV2 precursor into another may have resulted in the separation of ORF1 (Fig. 8B; scenario A). Retroelements like retrotransposons and EPRVs tend to integrate in a nested manner (SanMiguel *et al.*, 1996; Jakowitsch *et al.*, 1999), and in sugar beet, this was observed for several LTR retrotransposons as well (Weber and Schmidt, 2009; Wollrab *et al.*, 2012). The transduction of flanking DNA during this process presumably led to the acquisition of ORF3 between ORF1A and ORF1B followed by deletion of most of the integrated sequence, thus creating the same 5’-end in both components.

On the other hand, internal deletions may have happened first, forming two differently truncated beetEPRV2 entities (Fig. 8B; scenario B). One of these entities gained an additional ORF, possibly by the xenologous recombination (McClure, 2000) with a foreign pararetroviral sequence, as ORF3 bears no semblance to any other sequence in beet. Many viruses and repeats have modularly acquired domains and ORFs (Smyshlyaev *et al.*, 2013; Koonin *et al.*, 2015), some instances were also reported in beet (Heitkam *et al.*, 2014). Such an acquisition of ORFs likely contributed to the evolution of EPRVs, potentially leading to the speciation into virus families (McClure, 2000). In beet, we detected the resulting beetEPRV2 components A and B as both independent and combined insertions. We assume that recombination with a foreign pararetrovirus, as well as between the two beetEPRV2 components may have been enabled by similarities between their terminal repeats. One way or the other, the shared sequences between both beetEPRV2 components may have facilitated their mobilization as separate entities.

### Targeted integration or toleration – beetEPRVs accumulate in repetitive environments

Most of the detected beetEPRV members were embedded in repeat-rich environments as evidenced by sequence analysis (Fig. 4) and FISH (Fig. 7). On a nucleotide-level, the flanking regions (usually several hundred bases) of beetEPRV1 and beetEPRV3 contained a high amount of TA dinucleotides present as simple repeats. EPRVs with adjacent (TA)n repeats were detected in various plant species (*Oryza* sp.: Kunii *et al.*, 2004; and Liu *et al.*, 2012; various species: Geering *et al.*, 2014), indicating a potential pararetrovirus integration preference and retention. Besides the weaker electrostatic attraction between A and T nucleotides with only two hydrogen bonds, TA dinucleotide-rich sites can form secondary structures that disturb the DNA replication and thereby lead to instability within the chromosomes (Dillon *et al.*, 2013). Hence, the frequency of double strand breaks increases, as does the likelihood of a pararetroviral insertion during DNA repair. Consequently, beetEPRV integration may critically depend on the TA frequency in a potential genomic target.

On a macro-scale, 3-16 % of the beetEPRV sequences were flanked by satellite DNA arrays (Fig. 4), i.e. the major centromeric pBV and the intercalary pEV families (Schmidt and Metzlaff, 1991; Zakrzewski *et al.*, 2013; Schmidt *et al.*, 1991). This was also corroborated by our beetEPRV3 FISH (Fig. 7). The high TA-content of the centromeric satDNA in beet (59-69 %; Schmidt and Metzlaff, 1991) has presumably provided a suitable target for beetEPRV insertions. Although, centromeres are generally reduced in meiotic recombination (Bennetzen, 2000), centromeric repetitive DNA still evolves by unequal recombinatorial exchange (Ma and Jackson, 2006; Talbert and Henikoff, 2010). Thus, mitotic and meiotic DNA breaks may have been filled by beetEPRVs as observed for other retroelements (reviewed by Schubert and Vu, 2016).

A potential accumulation in DNA breaks may also explain the beetEPRV tendency to form clusters. Generally, clustering may result from the simultaneous involvement of several EPRV copies (e.g. EPRV concatemers) in a single break repair event, from a nested integration, or from the recombination of episomal viral genomes with integrated forms (Hohn *et al.*, 2008). EPRV clusters were observed in various plant genomes (e.g. tobacco, Lockhart *et al.*, 2000; petunia, Richert-Pöggeler *et al.*, 2003; rice, Liu *et al.*, 2012; Citrinae sp., Yu *et al.*, 2019). In *B. vulgaris*, EPRV accumulations are scattered throughout the entire genome and usually only contain beetEPRV members of the same family. This may be explained by rolling circle amplification or integration through recombination (e.g. Jakowitsch *et al.*, 1999; Kirik *et al.*, 2000; Gayral *et al.*, 2008), which depends on sequence similarity that is given within the respective beetEPRV family.

The specific distribution of beetEPRV3 on particular chromosome pairs and its association with specific host genome sequences may result from selective integration, retention, or removal, potentially also dictated by the three-dimensional structure of the genome (Bousios *et al.*, 2020). Regarding a selective integration, EPRVs do not encode an integrase and are, to our knowledge, unable to recognize specific target sequences. Nevertheless, EPRVs may be mobilized together with adjacent transposable elements, thus hitch-hiking to a location preferred by the transposable element (Staginnus and Richert-Pöggeler, 2006). Regarding removal and retention, we have observed EPRV depletion in the euchromatin: As euchromatic EPRVs are more likely to reduce plant fitness by interfering with gene expression, there would be a negative selective pressure in favor of EPRV removal. EPRVs in the heterochromatin, on the other hand, would likely be suppressed by their inactive chromatin environment, limiting detrimental effects and allowing EPRV retention (Hohn *et al.*, 2008). The survival of repeats in the heterochromatin, which form so-called ‘safe havens’ has been described for a number of repeats, mostly retrotransposons (Boeke and Devine, 1998; Gao *et al.*, 2008). Thus, the accumulation of beetEPRVs in repeat-rich environments is presumably the result of active selective targeting or passive retention in the heterochromatin or a combination of both.

### beetEPRV endurance – low level transcription despite silencing through RNA interference

Similar to LTR retrotransposons, the spreading of EPRVs requires reverse transcription of the RNA intermediate derived from the endogenous sequence. The arising episomal pararetroviruses are potentially virulent and can cause diseases such as banana streaks induced by the badnavirus BSV (Harper *et al.*, 1999), or vein clearing in tobacco induced by the solendovirus TVCV (Lockhart *et al.*, 2000) and in petunia induced by the petuvirus PVCV (Richert-Pöggeler *et al.*, 2003), respectively. For both LTR retrotransposons and EPRVs, the transcriptional activity depends on the required sequence motifs in the genomic copy to facilitate transcription by the host RNA polymerase, but may be counteracted by the host through epigenetic regulatory mechanisms (Ghoshal and Sanfaçon, 2015).

The potentially active endogenous PVCV sequence is characterized by flanking quasi-long tandem repeats (QTRs) that are assumed to have an LTR-like function facilitating its transcription. The QTRs comprise promoter and polyadenylation sequences and usually separate two adjacent PVCV sequences from another (Richert-Pöggeler *et al.*, 2003). Thus, transcription may start at the promoter of the upstream copy and terminate at the polyadenylation site of the consecutive one. PVCV also harbors a polypurine tract (5’-TTGATAAAAGAAAGGGGT-3’; Richert-Pöggeler and Sherpherd, 1997) that is supposed to function as primer binding site for plus-strand synthesis during reverse transcription. The TRs that we detected at the ends of all three beetEPRV families do not have an upstream polypurine tract and in case of beetEPRV3 also lack the poly(A) region that might act as polyadenylation signal. Thus, beetEPRVs do not contain canonical QTRs as described for PVCV and hence beetEPRV transcription and reverse transcription might be impaired. Nevertheless, we detected beetEPRV sequences in the published cDNA library of beet (genebank accession SRX674050), implying that basal EPRV transcription takes place.

As we detected small RNAs (smRNAs) matching the beetEPRV consensus sequences (Fig. 5) and infrequent cutting with methylation-sensitive restriction enzymes (Fig. 6), we assume that beetEPRV transcription is under epigenetic control. The beetEPRV-matching smRNAs target both coding and non-coding domains. However, the highest peaks were found in beetEPRV3 ORF2 and the region homologous to ORF2 of beetEPRV1, highlighting that the presumed FEV-specific protein encoded by ORF2 (Geering *et al.*, 2014) may be important for beetEPRV replication.

The three beetEPRV families are specifically targeted by smRNAs of variable lengths between 18 and 30 nt, indicating that beetEPRVs could be silenced by both transcriptional (TGS) and posttranscriptional gene silencing (PTGS). TGS is based on RNA-dependent DNA methylation which is mostly mediated by 24 nt smRNAs, while the hallmarks of PTGS are predominantly 21-22 nt smRNAs that lead to mRNA degradation and translation suppression of the targeted sequence (Ghoshal and Sanfaçon, 2015; Rosa *et al.*, 2018). We found that the beetEPRV families differ in the amount of matching smRNAs, as well as in their classification as either TGS- or PTGS-mediating smRNAs.

For beetEPRV2 we found low, mainly TGS involved smRNAs. The TGS pathway is also predominant for transposable element silencing in sugar beet (Zakrzewski *et al.*, 2013; Dohm *et al.*, 2014) characterized by high DNA methylation levels and contributing to the formation of large heterochromatic regions (Weber and Schmidt, 2009; Weber *et al.*, 2010; Zakrzewski *et al.*, 2011; Zakrzewski *et al.*, 2017). For EPRVs in particular, this silencing route was also observed in other host plants, such as petunia (Richert-Pöggeler *et al.*, 2003), rice (Kunii *et al.*, 2004), tomato (Staginnus *et al.*, 2007), and rapeseed (Omae *et al.*, 2020). The *Fritillaria imperialis*-specific EPRV (FriEPRV) was also found to be targeted by mostly TGS inducing 24 nt smRNAs (Becher *et al.*, 2014) and similar to beetEPRV2, no complete EPRV sequence could be identified for FriEPRV. However, as recombination of EPRV fragments have been observed to lead to the generation of complete and active viral genomes (Chabannes *et al.*, 2013), simple sequence rearrangements that do not loose vital parts of the virus genome as in the case of beetEPRV2 might not be sufficient to avoid viral activation.

In stark contrast, for beetEPRV1 and beetEPRV3, we detected mostly PTGS-specific lengths, supporting our assumption of a basal beetEPRV transcription, possibly by read-through mechanisms. Therefore, silencing has to take place at the posttranscriptional level to be effective. Strikingly, for beetEPRV3, the number of smRNAs is 13 to 36 times as high as for beetEPRV1 and beetEPRV2, respectively. This goes along with the observation that beetEPRV3 members are highly conserved and not fragmented, whereas beetEPRV1 and beetEPRV2 are usually marked by truncation and reshuffling. Consequently, we assume that (1) smRNA silencing is used to suppress a possible activation of beetEPRV3, and (2) that beetEPRV3 may be capable to become active, replicate and could be potentially infectious, if not hindered by the host.

As we detected beetEPRV1 and beetEPRV3 adjacent to the centromeric satellite DNA family pBV, and *in situ* hybridization also revealed beetEPRV3 clusters close to the centromeres of four chromosomes, the epigenetic centromere maintenance processes may also play a role in beetEPRV regulation. To initiate and maintain the centromere by incorporation of the centromere-specific histone H3 (CENH3) into the nucleosomes, transcription of centromeric DNA is required (Jiang *et al.*, 2003): During transcription, chromatin is disrupted and nucleosomes are destabilized and removed, providing an ideal opportunity for histone replacement. Any centromere-embedded transcription unit that comprises a promoter, including transposable elements, can initiate transcription.

This may explain why we found transcriptionally active beetEPRV1 and beetEPRV3 members and speculate that these are the copies associated with the centromeric repeats. Further, integration near transcriptionally active regions, such as the ribosomal genes, may also result in an unavoidable transcription of the adjacent pararetroviral sequences and subsequent silencing through PTGS. It is not uncommon that other repetitive elements are indeed adjacent or even embedded in the 18S-5.8S-25S rDNA (Balint-Kurti *et al.*, 2000; Jo *et al.*, 2009; Weber *et al.*, 2013). In case of beetEPRV3, we noticed an association with the 18S-5.8S-25S rRNA genes, localized distally on chromosome 1 (Schmidt *et al.*, 1994; Dechyeva and Schmidt, 2006) that would allow such transcription.

As our smRNA quantification relies on the mapping to beetEPRV consensus sequences, family members with a deviating sequence may potentially differ in smRNA coverage. Nevertheless, we can confidently state a high abundance of beetEPRV-derived smRNAs. These potentially contribute to the sugar beet resistance to infection with exogenous viruses of related sequences (Huang and Li, 2018), i.e. beetEPRV-derived smRNAs may play a role in the suppression of infectious, potentially pathogenic EPRVs. Therefore, beetEPRVs may represent a beneficial component of the host’s genome.

### Conclusion

The three beetEPRV families in sugar beet are characterized by differences in their structure, their genomic context and the ways in which they are silenced. Nevertheless, their common affiliation to the *Florendovirus* genus demonstrates their close relationship. Based on the low complexity organization of beetEPRV1 members and the high sequence similarity between beetEPRV2 and beetEPRV3, we postulate that the three beetEPRV families originated from at least two independent integration events 13.4 to 7.2 mya, with subsequent diversification of beetEPRV2 from beetEPRV3. During this process, the beetEPRV2 precursor probably underwent structure-changing mechanisms leading to its bipartite nature that we observe today.

BeetEPRV integration into the beet genome seems to be facilitated by a high AT amount in the genomic context. The accumulation of beetEPRV sequences in clusters points to active selective targeting and/or passive retention in primary heterochromatic regions while selective beetEPRV removal from the euchromatin of the host genome may also play a role. The observed embedment of beetEPRV sequences in the deep AT-rich heterochromatin of the host may ensure that the pararetroviral sequences remain inaccessible for several DNA-binding factors, thus reinforcing their silencing by the epigenetic control machinery. While beetEPRV2 is mostly targeted by smRNAs inducing DNA methylation, beetEPRV1 and beetEPRV3 show a high coverage with smRNAs inducing mRNA degradation and translation suppression, presumably as a result of basal transcription of the latter two beetEPRV families.

Taken together, the sugar beet host employs three strategies to shut down the beetEPRV copies, thus preventing re-infection: heterochromatic burial, epigenetic silencing, and structural disassembly. As a result, EPRVs in beet provide an example for the complete assimilation and inactivation of a plant virus in the host genome.

## Supporting information

Supplemental Data 1 (Fasta)

Supplemental Data 2 (Fasta)

Supplemental Data 3 (Fasta)

Supplemental Data 4 (Fasta)

Supplemental Data 5 (Fasta)

## SUPPLEMENTARY DATA

Additional supporting information may be found in the online version of this article and consists of the following. Figure S1: Specificity of EPRV nHMMs for the MP and RT domains and considerations for the parameter choice. Figure S2: Schematic representation of beetEPRV sequence variants and their occurrence in the EL10 assembly and on the PacBio long (SMRT) reads respectively. Figure S3: Localization of beetEPRV3 along mitotic prometaphase and metaphase chromosomes of *B. vulgaris*. Table S1: EPRV reference sequences used for phylogenetic analyses and their sources. Table S2: Primer sequences for the amplification of beetEPRV3 specific probes. Table S3: FEV-specific amino acids in the RT and MP revealed by the alignment of 16 EPRV sequences (see Fig. 2; Table S1).

Additional sequence data for beetEPRV reference sequences is accessible under http://doi.org/10.5281/zenodo.3888270 and comprises the following. Data S1: BeetEPRV reference sequences in fasta format. Data S2: Multiple sequence alignment of 27 beetEPRV1 sequences in fasta format. Data S3: Multiple sequence alignment of 42 beetEPRV2-A sequences in fasta format. Data S4: Multiple sequence alignment of 23 beetEPRV2-B sequences in fasta format. Data S5: Multiple sequence alignment of 11 beetEPRV3 sequences in fasta format.

## ACKNOWLEDGEMENTS

We want to dedicate this manuscript to Prof. Dr. rer. nat. Thomas Schmidt, our group leader, PhD supervisor, mentor and colleague. The identification of beetEPRVs have been on the mind of Thomas and his group and collaborators for a long time. A number of people have helped us to get to this point and we thank them for their initial work. We thank Pat Heslop-Harrison for advice and stimulating discussions.

We wish to thank the scientists involved in beet genome sequencing for providing early and pre-publication access to their data, thereby speeding up our research process. Furthermore, we acknowledge the genebank at the IPK Gatersleben for providing plant seeds.

