## Supplemental Tables and Figures for "Broken, silent, and in hiding: Tamed endogenous pararetroviruses escape elimination from the genome of sugar beet (*Beta vulgaris*)"

#### SUPPLEMENTARY TABLES

**Table S1:** EPRV reference sequences used for phylogenetic analyses and their sources

| genus | abbreviation | full name | source |
|---|---|---|---|
| *Caulimovirus* | CaMV | Cauliflower mosaic virus | genebank accession NC_001497[1] |
| | FMV | Figwort mosaic virus | genebank accession NC_003554[1] |
| *Soymovirus* | SbCMV | Soybean chlorotic mottle virus | genebank accession NC_001739[1] |
| *Badnavirus* | BSVAV | Banana streak VA virus | genebank accession AY750155[1] |
| | ComYMV | *Commelina* yellow mottle virus | genebank accession NC_001343[1] |
| *Tungrovirus* | RTBV | Rice tungro bacilliform virus | genebank accession NC_001914[1] |
| *Cavemovirus* | CSVMV | Cassava vein mosaic virus | genebank accession NC_001648[1] |
| *Solendovirus* | TVCV | Tobacco vein clearing virus | genebank accession NC_003378[1] |
| *Petuvirus* | PVCV | Petunia vein clearing virus | genebank accession NC_001839[1] |
| | FriEPRV | *Fritillaria imperialis* EPRV | https://onlinelibrary.wiley.com/doi/full/10.1111/tpj.12673[2] |
| *Florendovirus* | FEV *Atrich*BV | *Amborella trichopoda* B virus | https://www.nature.com/articles/ncomms6269#Sec18[3] |
| | FEV *Gmax*V | *Glycine max* virus | |
| | FEV *Ljap*AV | *Lotus japonicus* A virus | |
| 1 | Llorens *et al.* 2009, 2011 (Gypsy Database) | | |
| 2 | Becher *et al.* 2014 (Supplementary data S1) | | |
| 3 | Geering *et al.* 2014 (Supplementary data 1) | | |

**Table S2:** Primer sequences for the amplification of beetEPRV3-specific probes

| Primer | Sequence | Length of the predicted amplicon | Tm [°C] |
|---|---|---|---|
| bEPRV3_RT_for | CTC TCC AAT GGA TTC GTT ACC C | 389 bp | 58 |
| bEPRV3_RT_rev | GAA AAG CTT CAT CTT TGG TGC AG | | 57 |
| bEPRV3_MP_for | GCA TCC AAA GAT ATG AGT CTC C | 403 bp | 56 |
| bEPRV3_MP_rev | TAC ACT TTG GAT TAA GCT GAG AG | | 55 |

**Table S3:** FEV-specific amino acids in the RT and MP revealed by the alignment of 16 EPRV sequences (see Fig. 2; Table S1)

| Protein | Position in alignment (see Fig. 2) | FEV-specific amino acid | | Different chemical property compared to other EPRV sequences |
|---|---|---|---|---|
| RT | 4 | L/M | Leucine/Methionine | hydrophobic |
| | 8 | C | Cysteine | n/a |
| | 60 | E | Glutamate | negatively charged |
| | 79 | W | Tryptophan | aromatic |
| | 80 | I | Isoleucine | n/a |
| | 88 | K | Lysine | positively charged |
| | 129 | F | Phenylalanine | aromatic |
| | 147 | E | Glutamate | negatively charged |
| | 151 | I | Isoleucine | aliphatic |
| | 154 | D/E | Aspartate/Glutamate | n/a |
| | 210 | T | Threonine | n/a |
| | 213 | R | Arginine | positively charged |
| | 217 | H | Histidine | aromatic; positively charged |
| MP | 44 | P | Proline | n/a |
| | 85 | P | Proline | n/a |
| | 89 | N/D | Asparagine/Aspartate | n/a |
| | 90 | C | Cysteine | n/a |

### SUPPLEMENTARY FIGURES

#### EPRV identification

**A** nHMMs evaluation using a custom library of EPRVs and retrotransposons

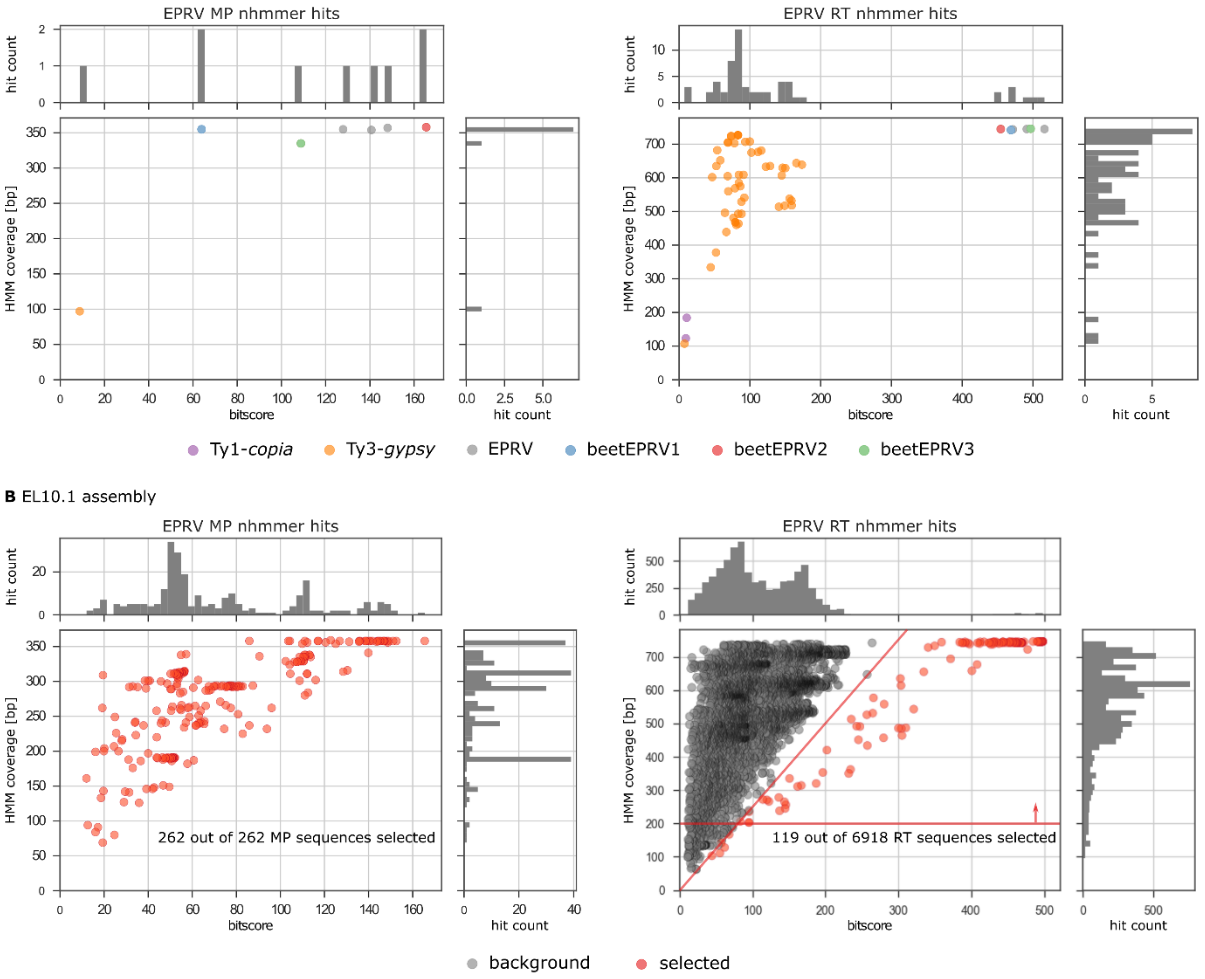


Figure S1: Specificity of EPRV nucleotide Hidden Markov Models (nHMMs) for the movement protein (MP) and reverse transcriptase (RT) domains and considerations for the parameter choice. For each of the analyses with an nHMM, we represent all hits as a dot in a scatter plot, based on the hit's bitscore and the length of the respective nHMM covered by the hit. In this representation, optimal hits are expected in the upper right corner of the scatter plot with high bitscore and high hmm coverage.

(A) The reference dataset, containing full-length sequences of 78 Ty1-copia, 19 Ty3-gypsy, and six EPRVs (Llorens et al., 2011) was queried independently with nHMMs for RT and MP. For the MP nHMM (left), except for one false positive Ty3-gypsy hit with a very low bitscore (orange), only EPRV sequences have been detected. For the RT nHMM, considerable cross-detection of Ty3-gypsy RTs (orange) was observed, however all with bit scores below 200. All EPRV RTs were detected with bitscores over 400.

(B) Hits from nHMM-based analysis on the EL10.1 reference assembly for beet (Funk et al., 2018) are plotted similarly to (A). For the downstream analysis, all MP hits (left) were included. For the RT (right), two cut-offs were defined: at least 200 bp coverage of the nHMM (red line) and with a quotient 1.5 ≤ (hmm coverage/bitscore) ≤ 2.5 (red dots). The latter enables the detection of shorter hits that still exhibit a high similarity to the nHMM.

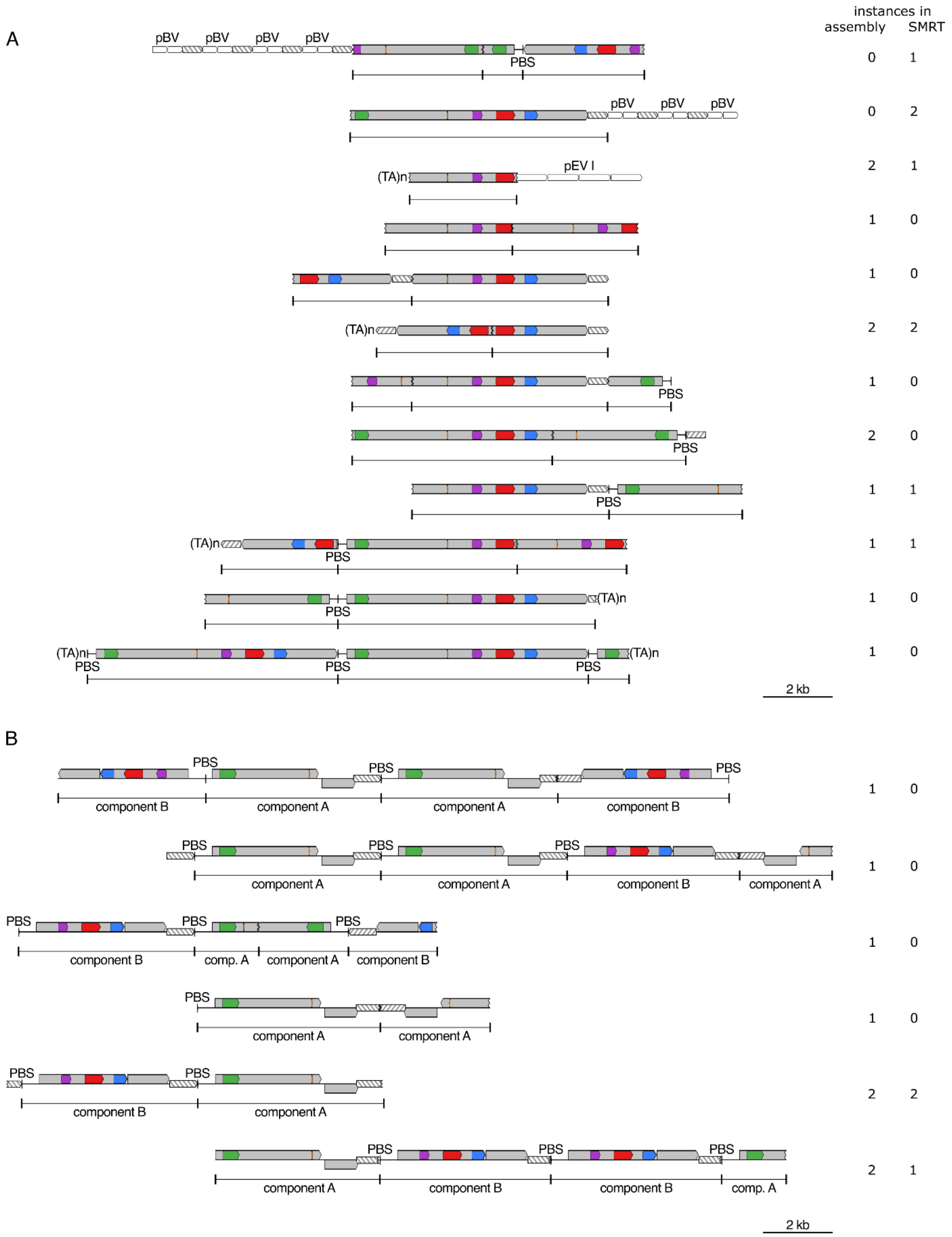

instances in
assembly SMRT
A
pBV pBV pBV pBV
PBS
0 1
pBV pBV pBV
0 2
(TA)n
pEV I
2 1
1 0
1 0
(TA)n
2 2
PBS
1 0
PBS
2 0
PBS
1 1
(TA)n
PBS
1 1
PBS
(TA)n
1 0
(TA)n
PBS PBS PBS
(TA)n
1 0
2 kb
B
PBS PBS PBS
component B component A component A component B
1 0
PBS PBS PBS
component A component A component B component A
1 0
PBS PBS PBS
component B comp. A component A component B
1 0
PBS
component A component A
1 0
PBS PBS
component B component A
2 2
PBS PBS PBS
component A component B component B comp. A
2 1
2 kb

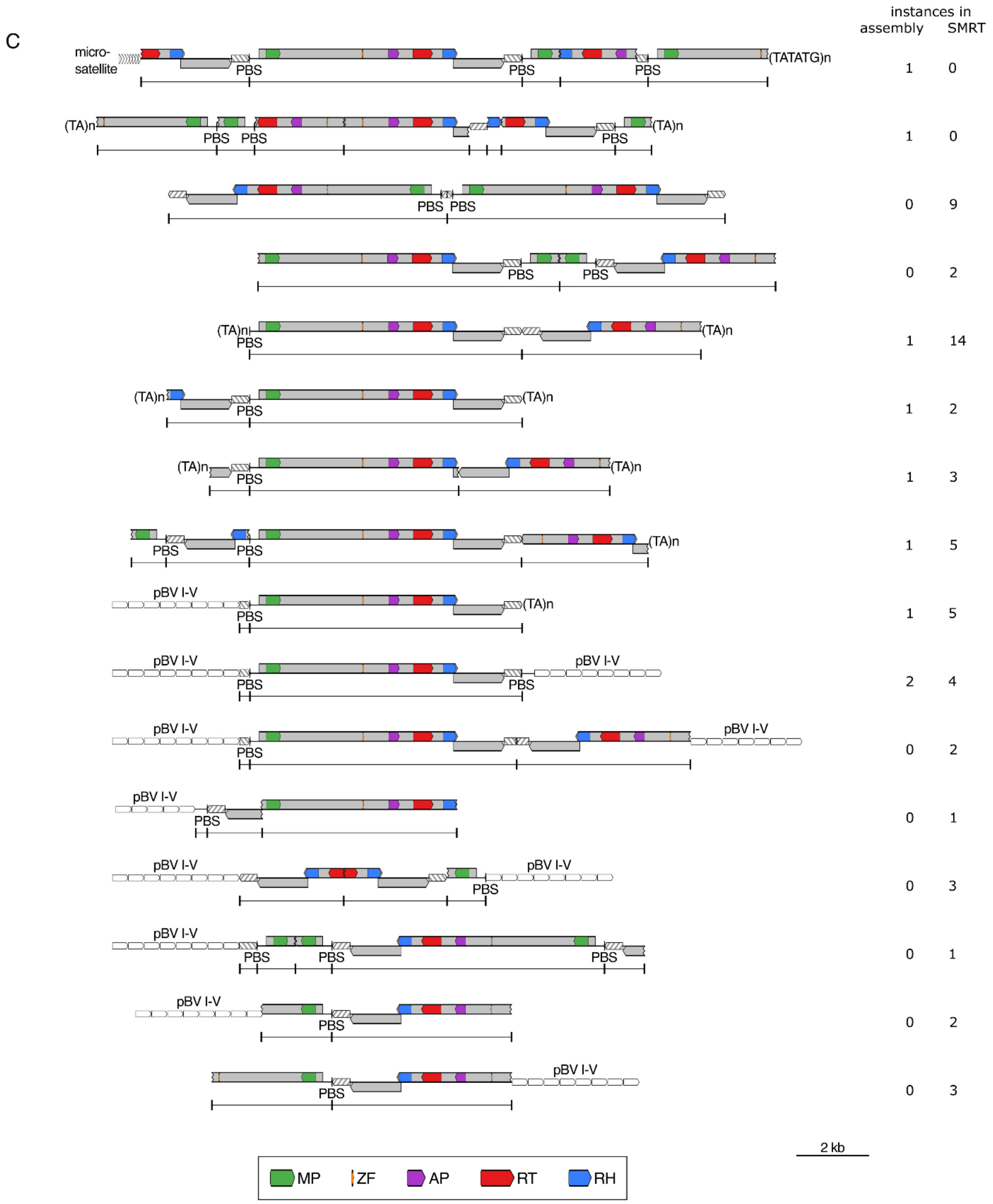


Figure S2: Schematic representation of beetEPRV structural variants and their occurrence in the EL10 assembly and on the PacBio long (SMRT) reads, respectively. Grey boxes: open reading frames; hatched boxes: terminal repeats; PBS = primer binding site; MP = movement protein; ZF = zinc finger in the coat protein; AP = aspartic protease; RT = reverse transcriptase; RH = RNase H1. Sequence variants are presented for beetEPRV1 (A), beetEPRV2 (B), and beetEPRV3 (C).

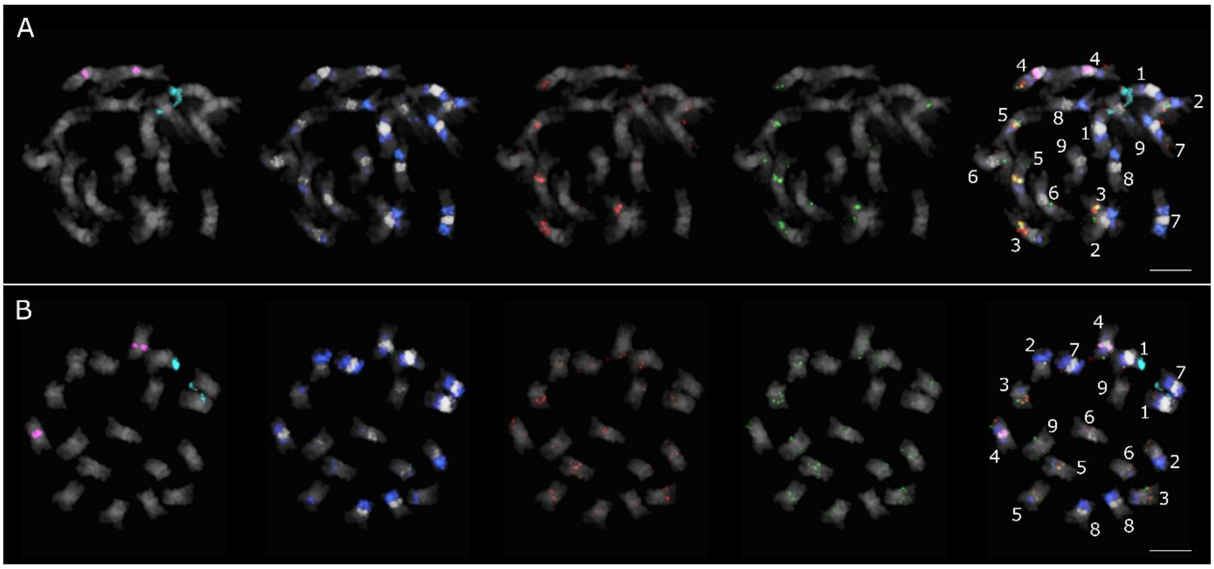


Figure S3: Localization of beetEPRV3 along mitotic prometaphase (A) and metaphase (B) chromosomes of B. vulgaris. DAPI-stained mitotic chromosomes of B. vulgaris are shown in grey. Multicolor FISH of 18S rDNA genes (turquoise), 5S rDNA genes (magenta), centromeric pBV I satellite (white), intercalary pEV I satellite (blue), beetEPRV3-RT (red), and beetEPRV3-MP (green). Information on probe labelling and detection can be found in the materials and methods section. Chromosomes were identified and numbers assigned following Paesold et al. (2012) and is based on the rRNA genes (pair one and four) and on the distribution and density of the satellites pBV I and pEV I.

Bars = 2 µm
